# Recombination within the *Cepaea nemoralis* supergene is confounded by incomplete penetrance and epistasis

**DOI:** 10.1101/472456

**Authors:** Daniel Ramos Gonzalez, Amaia Caro Aramendia, Angus Davison

**Affiliations:** School of Life Sciences, University of Nottingham, Nottingham NG7 2RD, United Kingdom; Universidad del País Vasco, Departamento de Zoología y Biología Celular Animal, Paseo Univ 7, Vitoria 01006, Spain

**Keywords:** *Cepaea*, colour, polymorphism, snail, supergene

## Abstract

Although the land snail *Cepaea nemoralis* is one of the most thoroughly investigated colour polymorphic species, there have been few recent studies on the inheritance of the shell traits. Previously, it has been shown that the shell polymorphism is controlled by a series of nine or more loci, of which five make a single ‘supergene’ containing tightly linked colour and banding loci and more loosely linked pigmentation, spread band and punctate loci. However, one limitation of earlier work was that putative instances of recombination between loci within the supergene were not easily verified. We therefore generated a new set of *C. nemoralis* crosses that segregate for colour, banding and pigmentation, and several other unlinked shell phenotype loci. The snails were genotyped using a set of RAD-seq loci that flank the supergene, and instances of recombination tested by comparing inferred supergene genotype against RAD-marker genotype. We found no evidence that suspected ‘recombinant’ individuals are recombinant within the supergene - point estimates of recombination between both colour/banding, and colour/pigmentation loci are zero, with upper limits of 0.8 and 1.8%. Incomplete penetrance and epistasis are a better explanation for the apparent ‘recombinant’ phenotypes. Overall, this work therefore provides a resource for fine mapping of the supergene and other major shell phenotype loci. It also shows that the architecture of the supergene may not be as previously supposed.

## Introduction

Historically, some of the most important animals in studying colour polymorphism have been the land snails *Cepaea nemoralis* and the sister taxon, *C. hortensis*, because it is straightforward to collect them and record the frequencies of the different morphs in different locations and habitats (Cain and Sheppard 1950; Cain and Sheppard 1952; Cain and Sheppard 1954; Jones et al. 1977). There is also the benefit that the major loci that determine the polymorphism show simple Mendelian inheritance (Cook 1967; Jones et al. 1977). However, while ongoing and long-term studies on these animals continue to provide compelling evidence for the fundamental role of natural selection in promoting and maintaining variation in natural populations, as well as the impact of modern-day habitat change (Cameron and Cook 2012; Cook 2017; Silvertown et al. 2011), the last research on the inheritance of the loci that determine the polymorphism date to the late 1960s. This is a problem because now that there is finally some progress towards identifying the genes involved (Kerkvliet et al. 2017; Mann and Jackson 2014; Richards et al. 2013), it is important that laboratory crosses are available, to validate prior knowledge on the inheritance and for use in fine mapping recombination break-points from a genome sequence.

Previous work has shown that the shell polymorphism is controlled by a series of nine or more loci, of which five or more make a single ‘supergene’, containing linked shell ground colour (C), banding loci (B), band/lip pigmentation (P/L), spread band (S) and punctate (or ‘interrupted’; I) loci. In most reports, colour and banding have been found to be tightly linked, with recombination typically towards the lower end of 0-2% (Cain et al. 1960; Cook 1967; Cook and King 1966). The exceptions are a study by Fisher and Diver (1934), which reported recombination of ~20% between C/B, and two crosses in Cain et al. (1960) which showed recombination of ~16%, also between C/B. Although there have been fewer studies, pigmentation, spread band and punctate are more loosely linked, showing rates of recombination between 3 and 15% (Cain et al. 1960; Cain et al. 1968; Cook 1967). The main other loci that make up the shell phenotype are various forms of band-suppressing loci, all unlinked to the supergene, including the mid-band locus, *U* (unifasciata), and another that suppresses the first two bands, *T* (trifasciata).

One unavoidable limitation of prior works was that putative instances of recombination between loci within the supergene could not be verified, except by breeding further generations of snails from the ‘recombinant’ offspring to confirm the underlying genotype. As this was rarely possible, in some studies it was recognised that incomplete penetrance might be an alternative explanation for the phenotype of recombinants. Chance arrangements of alleles at other loci might sometimes interact to prevent expression of a particular phenotype, causing individuals to appear as if they are ‘recombinant’ (Cook and King 1966).

To further understand the frequency of recombination within the supergene, and to generate further material for fine mapping, we made a new set of *C. nemoralis* crosses that segregate for several shell phenotype loci. The offspring were then genotyped using a set of linked RAD-seq loci that flank either side of the supergene (Richards et al. 2013), and instances of recombination confirmed or refuted by comparing inferred supergene genotype against RAD-marker genotype - the underlying idea is that individuals that show recombination within the supergene should also be recombinant by RAD-marker. Overall, we found that the phenotype of ‘recombinant’ individuals is better explained by incomplete penetrance and epistasis.

This work therefore provides a method to identify recombination events that either flank the supergene or are between loci within the supergene. The results also show that recombination within the supergene may be considerably rarer than supposed.

## Materials and methods

### The culture of *Cepaea*

Snails were fed a hydrated grass pellet, oat and chalk mix, supplemented with lettuce, as described previously (Davison 2000). Generally, large juvenile virgin snails were raised to adulthood in isolation and then introduced to a partner. Pairs of snails were then kept in tanks with ~4 cm soil until egg laying began. As *C. nemoralis* is a simultaneous hermaphrodite, offspring from both parents were used. Egg batches were isolated, and the offspring reared to adulthood under the same feeding regime, with the time from egg to adult being ~6 months. In most cases, adult offspring, or occasionally, large subadult snails, and parents were preserved frozen.

The majority of snails were raised to adulthood, and so the colour, banding, band pigmentation and lip colour phenotype was scored and then the shell genotype inferred. Overall, the complications in scoring some characters necessitated minor deviations from the scheme put forward by Cain (1988).

We have previously shown that colour variation is multimodal but continuously variable in natural populations, necessitating the use of quantitative methods to measure it (Davison et al. 2018). However, in simple crosses it is straightforward to bin the individuals into one of two types; quantitative measures are not necessary. Therefore, the shell ground colour phenotype was scored as either yellow (Y), pink (P) or brown (B), and where possible, the corresponding genotype inferred (dominance is *C^B^* > *C^P^* > *C^Y^*), including whether in coupling or repulsion phase with other loci.

Similarly, banding phenotype was generally scored as either unbanded (O; 00000), mid-banded (M; 00300) or having several bands (B; most frequently 12345, but all combinations except 00300) and the genotype at the banding locus was inferred (unbanded dominant; *B^O^* > *B^B^*). Several crosses also segregated for the midband locus, so the genotype at that locus was also inferred (mid-band dominant; *U*^3^ > *U*^−^). One cross segregated for two other potential band-suppressing loci, *T* (first two bands missing: 00345; T^345^ > *T*^−^) and another possible locus, which we called X (first band missing or very faint: 02345; X^2345^ > X^−^).

It is not clear from previous studies as to whether the lip pigmentation is a separate locus, or is instead allelomorphic with the band pigmentation locus. As a precaution the two were therefore treated separately. Thus, band pigmentation phenotype was scored as normal (N) or hyalozonate (H) and the corresponding genotype inferred (*P^N^* > *P^h^*). Hyalozonate shells typically have unpigmented bands and lip (see Discussion); discrete bands can still be recognised because the background colour is paler than the shell ground colour. In some crosses, the lip pigmentation phenotype was two distinct types, either normal (L), or white lip (A; albolabiate), and so the corresponding genotype was inferred (*L^L^* > *L^a^*). In other crosses, lip pigmentation showed quantitative variation and so was difficult to score.

The adults used in crosses 10, 11, 12 and 13 were derived from offspring of cross 9, so the shell genotype could be inferred with extra confidence. This was aided by full-sib inbreeding in producing crosses 10, 11 and 12, and another round of inbreeding to produce cross 13. One benefit of the inbreeding is that yellow snails in these crosses must be wholly homozygous for the majority of the chromosome that contains the supergene, and so we could be absolutely confident about supergene genotype.

For each cross, Mendelian segregation ratios were tested using chi-square goodness-of-fit tests.

### DNA extractions

Genomic DNA was extracted from frozen snail foot tissue as described previously (Richards et al. 2013). For most samples, slices of snail tissue were incubated at 65°C in extraction solution (3% CTAB, 100 mM Tris-HCl, pH 7.5, 25 mM EDTA, pH 8, 2 M NaCl) with 0.2 mg/mL proteinase K and 80 μg/mL RNase. Upon lysis, a chloroform extraction was performed, then three volumes of CTAB dilution solution added (1% CTAB, 50 mm Tris-HCl, pH 7.5, 10 mM EDTA, pH 8). Samples were mixed until a precipitate appeared, then the supernatant removed. The pellet was washed twice in 0.4 M NaCl in TE (0.4 M NaCl, 10 mM Tris-HCl, pH 7.5, 1 mm EDTA, pH 8), re-dissolved in 1.42 M NaCl in TE (1.42 M NaCl, 10 mM Tris-HCl, pH 7.5, 1 mM EDTA, pH 8), then precipitated in ethanol, centrifuged and dried. Latterly, a few samples were lysed in lysis buffer (10mM Tris, 0.1M EDTA, 0.5% SDS), then extracted using the standard phenol-chloroform protocol.

### RAD-marker genotyping of parents and offspring

For future mapping and the precise identification of the supergene, it would be useful to identify individuals that have are known to have a recombination break-point close to the supergene, or better still, between the component loci (as previously). We therefore genotyped a subset of the individuals in several of the crosses using custom assays derived from RAD-seq loci that flank either side of the supergene. For this, standard PCR was carried out either using Amplitaq Gold polymerase (Invitrogen), 1.5 mM MgCl2, using a cycle of 95°C for 10 min, followed by 35 cycles of 95°C for 30 s, 58°C for 30 s, and 72°C for 1 min or Clontech Advantage 2 PCR with an initial denaturation of 95°C for 1 min, followed by 35 cycles of 95°C for 15 s, 65°C for 1 min, 68°C for 1 min, and 72°C for 1 min. Primers and custom genotyping assays were based on the previously characterised RAD-seq loci (Richards et al. 2013), and varied according to the cross. The primers were RAD06F 5’-GCCTATCCGTCATTGTTGGT-3’ RAD06R 5’-GTCAAGGCTTGCTTCTTTGG-3’, RAD9F 5’-TTTCTCGGAACGACGGAGT-3’, RAD9R 5’-GGTCTCGTCAATGGCACTTT-3’, RAD11F 5’-AAGAAGCGTCCTTCTGGAAA-3’, RAD11R 5’-CACCTTCCCCATTCTTCAAA-3’. For each cross it was necessary to custom design an assay, so that segregation of markers could be observed in the offspring. The enzymes used with each assay and each cross are detailed in Supplementary Table 1.

## Results

### Segregation of Mendelian loci that determine shell phenotype

Shell colour, locus C, showed segregation in crosses 1 to 13 (Table 1; Supplementary Table 2), only deviating significantly from expected Mendelian segregation ratios in cross 12, with fewer yellow shells than expected. Crosses 1 to 6 and 8 showed segregation for the band presence/absence locus, B, with no deviations from expected Mendelian ratios. Crosses 6 and 9 to 13 showed segregating variation for the mid-band phenotype, coded by the unlinked *U* locus. The observed phenotype frequencies did not differ from the expected frequencies. Crosses 10 to 13 showed segregation for the pigmentation (hyalozonate; *P*) locus, with no deviations from expected Mendelian ratios.

**Table 1.**
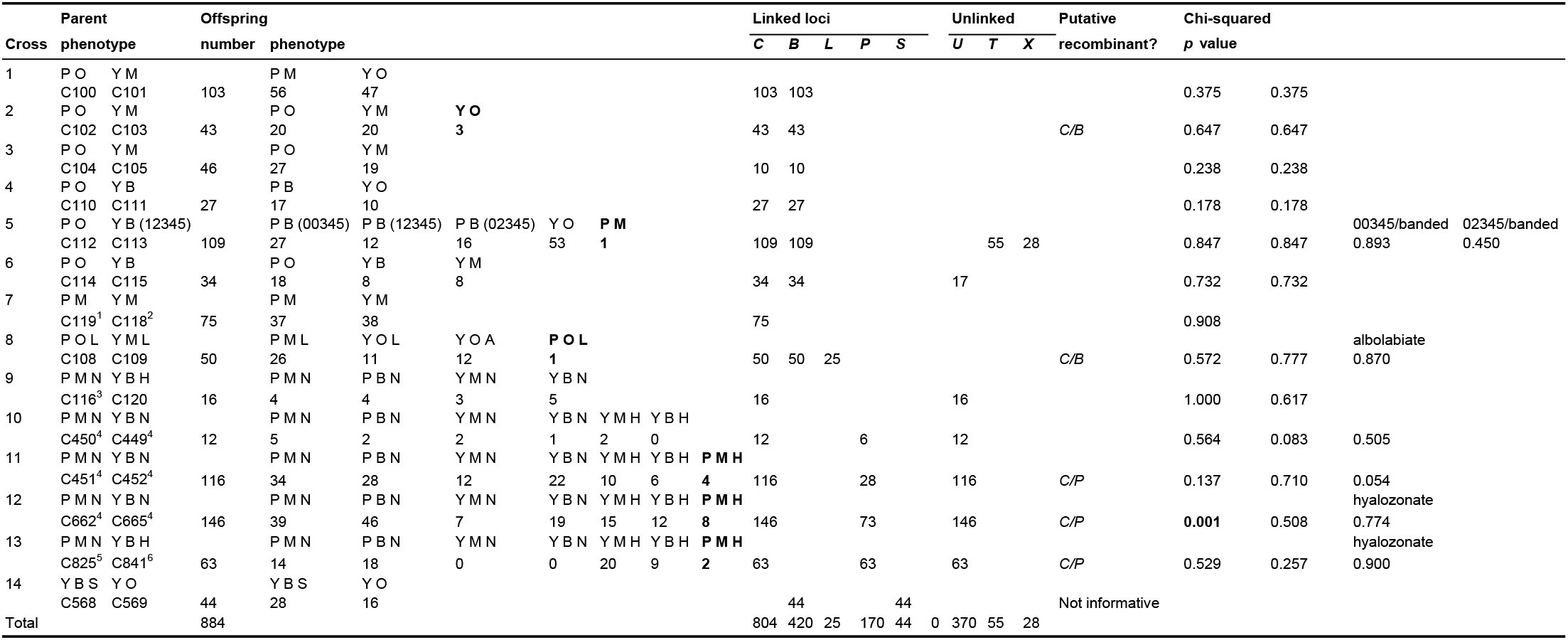
Summary of parent and offspring phenotypes from *C. nemoralis* crosses. Phenotypes that may be due to a recombination event in a parent are highlighted in bold. Inferred genotypes of offspring are detailed in Supplementary Table 1. Key: P pink, Y yellow, O unbanded, M mid-banded, B all other banding patterns; N normal band pigmentation; H hyalozonate banding (nearly always with white lip - see text); S spread banding; L normal lip pigmentation; A albolabiate (white lip). Cross 5 also showed segregation for another one or two band-suppressing loci, *T* and *X*, so the detailed banding notation is also shown.

Cross 8 showed segregation for the putative lip colour locus, *L*, with no deviations from expected Mendelian ratios, assuming that it is also part of the supergene and treating lip colour phenotypes was as either normal (N) or albolabiate lip (A). Offspring in several of the other crosses, especially 10 to 13, showed considerable and apparently continuous variation in lip colour. We therefore tried to score the lip phenotype in the conventional manner, having a phenotype as normal, pale, or albolabiate, and reconcile this with a knowledge of the parental phenotypes and genotypes (parents in crosses 8 and 9). No scheme that we devised fitted a simple Mendelian model. This fits with previous studies that have shown that incomplete dominance (Cain et al. 1968; Cook 2003). Unfortunately, it was also not possible to quantitatively measure the lip colour, because the coloured part of the lip was frequently too small and also on a curved surface.

Cross 14 showed segregating variation in the spread band phenotype. However, as one parent was homozygous for the dominant spread band allele *S^s^*, the cross was non-informative for recombination with other supergene loci.

Finally, cross 5 showed segregating variation for the locus that suppresses the first two bands, converting a five-banded snail (12345) to three-banded (00345). The offspring phenotype frequencies would be consistent with single locus, with *T^345^* dominant to *T*^−^, with both parents being heterozygote, except the problem is that this would require both parents to have a *T^345^* allele; apparently not possible because one of the parents is 12345. As five of the offspring with suppressed bands have 02345 phenotype and eleven a 0:345 (: = trace) phenotype, then the results are consistent with their being two band-supressing loci, one that causes the 00345 phenotype and another that causes the 02345/0:345 phenotype (see Supplementary Table 2 for inferred genotypes).

### Putative recombinants between colour, banding and lip and band pigmentation loci

Previously, the colour (P/Y) and banding (B/O) phenotype for six crosses (1, 4, 5, 6, 7, 8) and 398 offspring was reported, and the genotype of flanking RAD-seq loci reported for cross 1. In this new work, we raised a further 486 offspring from eight more crosses, and genotyped six further crosses using flanking RAD-seq loci. The combined data set of parent and offspring phenotypes, alongside inferred genotypes, is presented here together, summarised in Table 1, and presented in full in Supplementary Table 2. Offspring in several crosses produced snails with phenotypes that could be explained by a recombination event within the supergene of the heterozygous parent (Figure 1).

**Figure 1.**
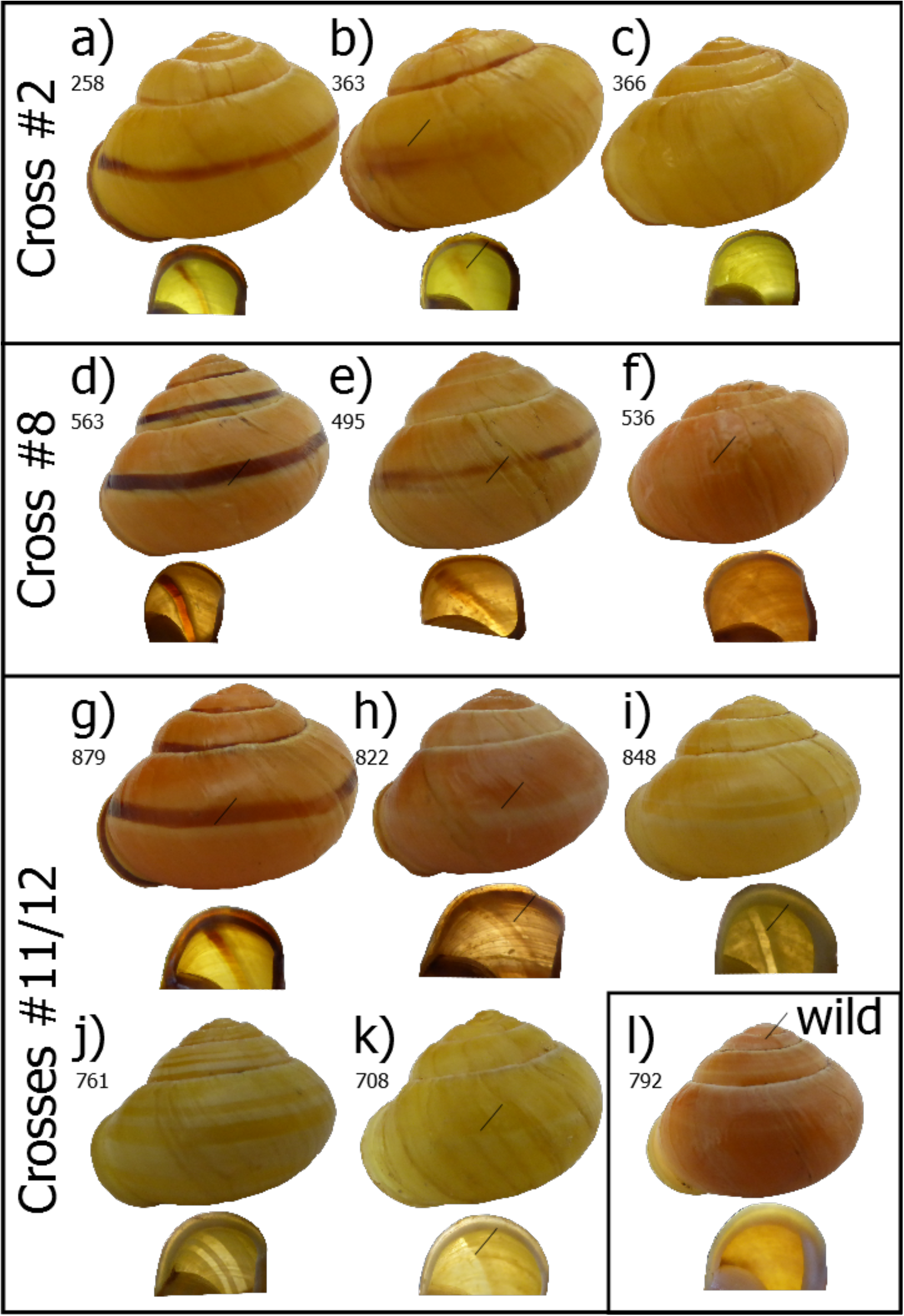
Shells of offspring from crosses, including putative recombinant individuals, and one wild collected individual. a) normal yellow mid-band, b) yellow, trace of banding, c) yellow, no band. An absence of banding suggests that individual 366 is a putative recombinant. d) normal pink mid-band, showing evidence of white “highlighting” of pigmented band e) pink, trace of banding, some highlighting f) pink, no band, very faint mark where band would be. An absence of banding suggests that individual 536 is a putative recombinant. g) normal pink mid-banded, showing evidence of white highlighting of pigmented band h) pink, no band, white highlighting i) yellow, mid-band hyalozonate, j) yellow, banded hyalozonate (02345) k) yellow, mid-band hyalozonate. An absence of dark pigment suggests that 822 is a putative recombinant; however, the shell has retained the white highlighting pigment. Hyalozonate shells generally lack both dark and light pigment, see i) and j); this is not always easily visible, see k). In a wild-collected pink hyalozonate, l) the lack of pigment is only just visible on the upper whorls, and not the last whorl or from the inside.

Cross 2 produced three yellow unbanded snails (e.g. Figure 1c), a phenotype that might be produced by recombination between the colour (*C*) and banding (*B*) loci; one of these individuals, a sub-adult with a damaged shell, has a very faint trace of a band. A few other snails in the same cross have much reduced banding (e.g. Figure 1b).

Cross 8 produced a single pink unbanded snail, a phenotype that is also best explained by recombination between the colour (*C*) and banding (*B*) loci (Figure 1f). Very few of the snails in this cross had reduced banding. This cross also segregated for the lip pigmentation locus, L. As the recombinant snail has a pigmented lip (see Figure 1f, lip image), then this cross in theory informs upon the order of loci within the supergene (but see below).

Cross 5 produced a single pink mid-banded snail. This phenotype is very difficult to explain by recombination, based on the known genotypes (Supplementary Table 1). As the pink colour of this snail is qualitatively different from the other pink-banded snails in this cross, the best explanation is that it is a likely a contaminant from another cross.

The remaining crosses produced offspring that suggest possible recombination between the colour *C* and band pigmentation *P* loci. Crosses 11, 12 and 13 produced several unbanded pink individuals, with pigmented lips (e.g. Figure 1h). These were initially scored as hyalozonate, because the mid-band was evident but not pigmented. However, closer inspection revealed that the banding phenotype of these shells is not the same as the yellow hyalozonate shells. Specifically, the unbanded pink individuals retain the white highlighting pigment of a normal shell, but lack the dark pigment (compare Figure 1h with 1g). This difference is especially evident when viewed from the underside: hyalozonate shells have cleared bands which are entirely lacking pigment whereas the white pigment of ‘unbanded’ snails shows a silhouette (compare Figure 1h with 1i, j). Not all of the hyalozonate shells show such a clear pattern (e.g. Figure 1k). For comparison, in a wild collected pink hyalozonate, the cleared bands are only evident on the upper whorls (Figure 1l); in another shell they are not evident at all.

### Genotyping of offspring using RAD-seq derived loci

Individual offspring from crosses 1, 2, 8, 9, 10, 11, 12, and 13 were genotyped using custom assays derived from RAD-seq loci that flank either side of the supergene, using RAD06/RAD11 on one side and RAD09 on the other side (Table 2; Supplementary Table 2). Unfortunately, the RAD-seq loci in cross 5 lacked polymorphism so no assay was possible. To confirm or refute individual recombination events, we inspected the genotype of the putative recombinant offspring. In theory, individuals for which we have inferred recombination within the supergene should also show recombination by one of the RAD-markers.

**Table 2.** Summary of RAD-seq marker genotyping, putative number of supergene recombinants and actual number. RAD06 and RAD11 flank one side of the colour and banding loci of the supergene; RAD09 flanks the other side. Full genotypes are in Supplementary Table 1.

| Cross<br>Snails and phenotypes |  |  |  |  | Genotypes |  |  | Recombinants between<br>supergene and: |  |  | Putative<br>supergene<br>recombinants | Actual<br>supergene<br>recombinants |
| --- | --- | --- | --- | --- | --- | --- | --- | --- | --- | --- | --- | --- |
|  |  |  |  |  | RAD11 | RAD06 | RAD09 | RAD11 | RAD06 | RAD09 |  |  |
| 1 | C100 | C101 | P O | Y M | 101 | 102 | 102 | 1 | 10 | 1 | 0 | 0 |
| 2 | C102 | C103 | P O | Y M | 38 |  | 43 | 0 |  | 2 | 3 | 0 |
| 8 | C108 | C109 | P O L | Y M L | 44 |  | 50 | 2 |  | 1 | 1 | 0 |
| 9 | C116 | C120 | P M N | Y B H |  | 16 | 16 |  | 2 | 0 | 0 | 0 |
| 10 | C450 | C449 | P M N | Y B N |  | 12 | 12 |  | 2 | 1 | 0 | 0 |
| 11 | C451 | C452 | P M N | Y B N | 103 |  | 104 |  | 4 | 2 | 4 | 1? |
| 12 | C662 | C665 | P M N | Y B N | 127 |  | 132 |  | 3 | 6 | 8 | 0 |
| 13 | C825 | C841 | P M N | Y B H | 44 |  | 49 |  | 4 | 1 | 3 | 0 |

None of the three individuals from cross 2 showed evidence of recombination from the flanking loci RAD11 and RAD09. Similarly, the single individual in cross 8 did not show evidence of recombination for the same RAD-seq loci. All eight putative recombinants in cross 12 and all three in cross 13 showed no evidence of recombination using RAD06 and RAD09; in cross 11, three individuals did not show evidence of recombination, with one single individual (804) showing recombination between RAD06 and the supergene and apparent recombination between the colour and pigmentation loci (C/P). If this were correct then it would inform the order of loci within the supergene. However, the phenotype of this snail is exactly the same as the other refuted recombinants (Figure 1h). It is most likely not a hyalozonate, and therefore a coincidence that it also show recombination between RAD06 and the supergene.

Thus, overall, while there is incomplete evidence in some cases, we were not able to confidently confirm any recombination events within the supergene. This puts the point estimate on recombination between *C* and *B* at 0/376 (<0.27%), between *C* and *P* at 0/170 (<0.60%) and between *C* and *L* at 0/25 (<4%). Therefore, the upper confidence limit for the mean rate of recombination, assuming that the probability of observing zero recombination events is still 5%, is 0.80% for *C*/*B*, 1.76% for *C*/*P*, and 12.0% for *C*/*L*. Of course, as there are no recombination events, then this work does not inform the order of loci within the supergene - the higher upper limits for *C*/*P* and *C*/*L* are due to smaller sample sizes.

## Discussion

For future mapping and the precise identification of the supergene, it would be useful to identify individuals that are known to have a recombination break-point close to, or within the supergene. In this study, we initially identified four putative recombination events between the colour and banding loci (*C*/*B*) and fourteen putative recombination events between the colour and pigmentation loci (*C*/*P*). This is as expected because historic studies have indicated that *C*/*B* are more tightly linked than *C*/*P*. We also used genotyping of RAD-seq loci that flank the supergene in several of the crosses to reveal individuals that show recombination between a linked RAD-seq marker and the supergene. These RAD-seq markers may therefore be used for future recombination break-point mapping. However, the same RAD-seq genotyping falsified the putative inferences for recombination *within* the supergene. This is also supported by a close analysis of the phenotype of the shells, including variable penetrance of the mid-band phenotype (Figure 1; crosses 2, 8, 11 to 13) and a comparison of true hyalozonate shells against shells that partially lack pigmentation (Figure 1; crosses 8, 11 to 13).

Therefore, in contrast to previous studies that have reported rates of recombination between *C*/*B* of 0-2% and between *C*/*B* and the pigmentation locus, P, of 3-15%, we found zero recombinants, putting upper limits on the rate of recombination at 0.8% and 1.8%, respectively. This does not mean that previous inferences of recombination within the supergene were incorrect. However, as the ‘recombinants’ in this study are better explained by other means, then we conclude that there is an absence of modern-day evidence for recombination within the *C. nemoralis* supergene. The structure of the supergene may not be as has previously been supposed.

### Incomplete penetrance and epistasis

Four individuals in crosses 2 and 8 were initially identified as putative recombinants, because they lacked the mid-band, or had only very faint traces of a band. If these snails had been true recombinant individuals then they would most likely be homozygous for colour and heterozygous for banding (genotype *B^0^B^b^*). Incomplete penetrance of the dominant *B^0^* allele could mean that some individuals show evidence of banding (e.g. Figure 1b, e). Instead, the genotyping shows that these individuals are not recombinant, and so must be genetically homozygous for colour and banding (*B^b^B^b^*). Therefore, the best explanation for their phenotype is that other loci are interacting epistatically to prevent full penetrance of homozygous banding alleles.

Similarly, up to fourteen individuals were identified that were putative recombinants between the colour and pigmentation loci. If they had been true recombinants, then they would most likely be heterozygous for colour and homozygous for pigmentation (*C^P^C^Y^P^H^P^H^*). However, close analysis of the phenotype and the genotyping together show that they are not recombinants and therefore, more likely they were heterozygous for both colour and pigmentation (*C^P^C^Y^P^N^P^H^*). These same snails are homozygous for the banding locus (*B^b^B^b^*), but segregate for mid-band phenotype (*U^3^U*^−^, *U*^−^*U*^−^); the putative recombinants were always mid-banded (*U*^3^*U*^−^), rather than fully banded (*U*^−^*U*^−^). The same explanation may apply, as above. Other loci sometimes interact to prevent full penetrance of the mid-banded phenotype.

The observation that the absence of a mid-band does not always have a simple genetic basis may shed some light on previous findings. For example, both Fisher and Diver (1934) and Cain et al. (1960) reported individual crosses that showed elevated rates of recombination between the *C*/*B* loci.

Cain et al. (1960) reported two crosses derived from the same mother that showed seven colour/banding recombinants in 43 snails, for which six were pink mid-banded snails. The expectation could be the reverse to the above, that recombinants should be homozygote for the banding locus (*C^P^C^Y^B^b^B^b^*) and non-recombinants heterozygous (*C^P^C^Y^B^0^B^b^*). Incomplete dominance of the band suppressing allele (*B^0^*), or else epistatic interactions with other loci, may be an alternative explanation for these ‘recombinant’ individuals.

Similarly, Fisher and Diver (1934) described unexpectedly high recombination (20%) between colour and banding in one cross. Unfortunately, they did not report whether the snails used were mid-banded or not. However, in their specific case, doubt has been raised as to whether the individuals used were virgins before paired together (Cain et al. 1960; Ford 1971; Lamotte 1954). In researching this work, we were fortunate to find copies of letters between Fisher and Diver in the archive of Bryan Clarke (Supplementary Material). In letters from April/May 1934 that describe the preparation of the correspondence that was published in Nature in June 1934, there is clear admission that the snails used in the crosses were adult and not virgin. The authors partly acknowledge that this may be a problem. Referring to possible previous matings (“experience”), Fisher writes that “on fairly strong ground, which is not weakened by previous experience, but is not absolutely critical”. According to our interpretation, Fisher also discounts previous matings as being a problem, because the offspring ratios approximate to what you would expect with limited recombination. There are therefore perhaps two errors, which together invalidate the conclusions of that correspondence.

Epistasis could also explain other earlier data on recombination between the *C*/*B* loci and the pigmentation locus. For example, in our study, we initially scored some pink individuals as hyalozonate, even though they had a lightly pigmented lip, only later realising our error. Other authors may have made the same mistake. Unfortunately, it is difficult to be certain from the previous literature whether pink hyalozonate recombinant individuals had an unpigmented lip, though if this was not the case then might it have been explicitly noted (e.g. p404 in Cook 1967)? However, just as there is epistasis so that brown shelled individuals are less likely to be banded, then it is reasonable to suppose that pink hyalozonates more rarely have a wholly unpigmented lip.

Evidently, further crosses are required, especially with respect to the other major loci, especially pigmentation, spread band and punctate loci in the supergene, and the various band-modifying genes. It is possible that some of these phenotypes, especially those for band-modification (Wolda 1969), are under multi-factorial control and/or dependent upon genetic background.

### Future progress

Overall, this work provides a resource for fine mapping of the supergene, and the other major shell phenotype loci. On the one hand, we have shown that phenocopies may be a problem in using the shell phenotype alone to detect recombination events within the supergene. On the other hand, the genotyping methods that we have introduced enable a means to avoid this problem. Jones *et al*. (1977) (in)famously questioned whether understanding polymorphism in *Cepaea* is “a problem with too many solutions?” The intention of that work was to emphasise the perfect case study provided by *Cepaea*, although that was not the effect. We hope that these crosses may soon be used with new long-read DNA sequencing methods to assemble the *C. nemoralis* genome and to identify the supergene. Then, perhaps polymorphism in *Cepaea* may instead be considered “a solution to many problems.”

### Data archiving

All relevant data is in the tables and Supplementary Material.

## Acknowledgements

This work was mainly funded by the University of Nottingham, a BBSRC studentship to DRG, a PhD and a visiting fellowship awarded by the Dept. of Education, Universities and Research of the Basque Government (Ref. PRE_2015_2_0191; Ref. EP_2015_1_33) to AC. Thanks to both Sheila Keeble and Julie Rodgers for help with the care of snails, and to Laurence Cook for comments on the manuscript, and to Anne Clarke and the University of Nottingham for access to the archive of Professor Bryan Clarke.

## Electronic supplementary material

Supplementary Tables 1 and 2. Supplementary information: scanned correspondence between Ronald A. Fisher and Captain Cyril Diver.

---

**Supplementary Table 1.**
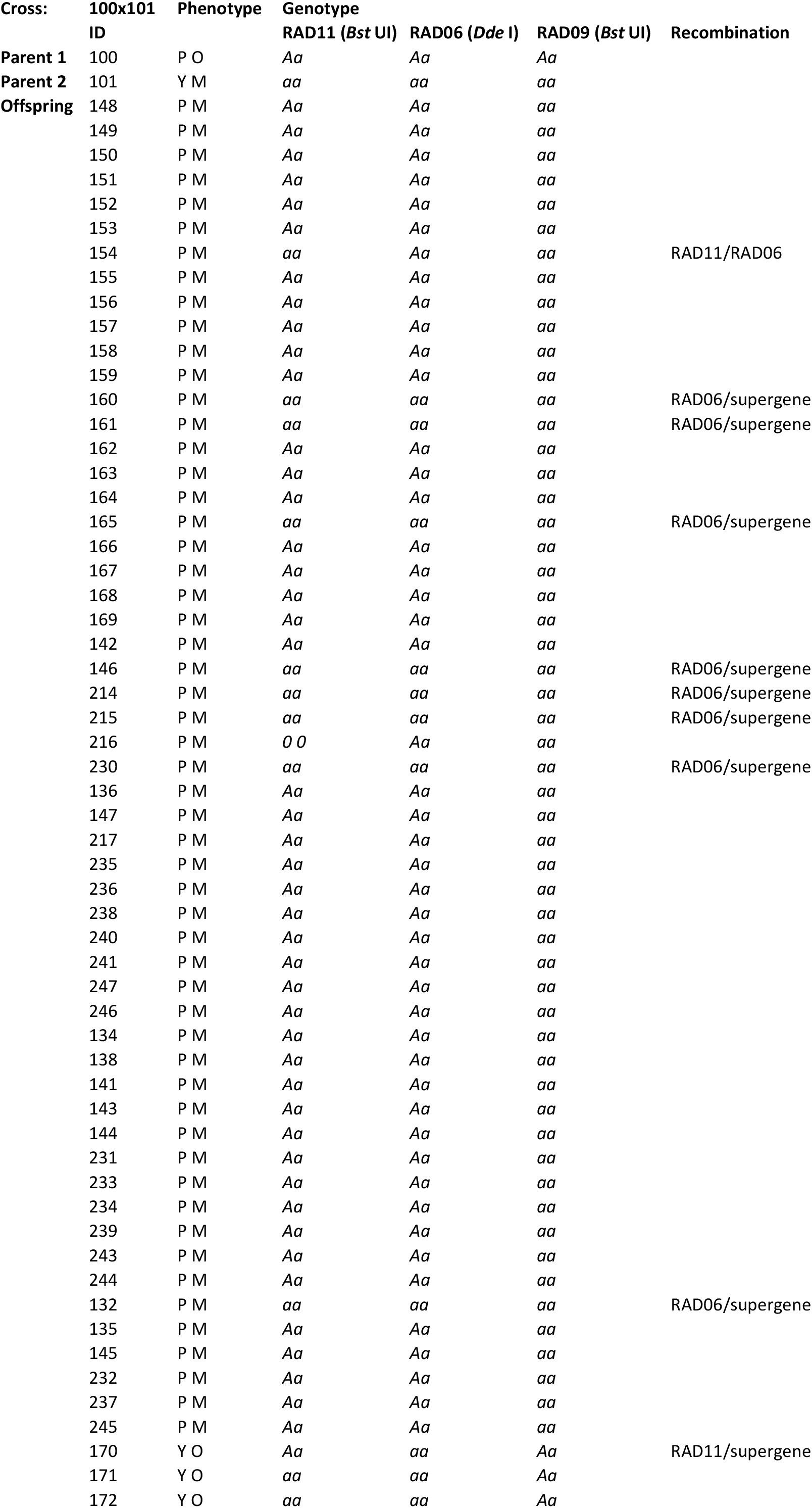
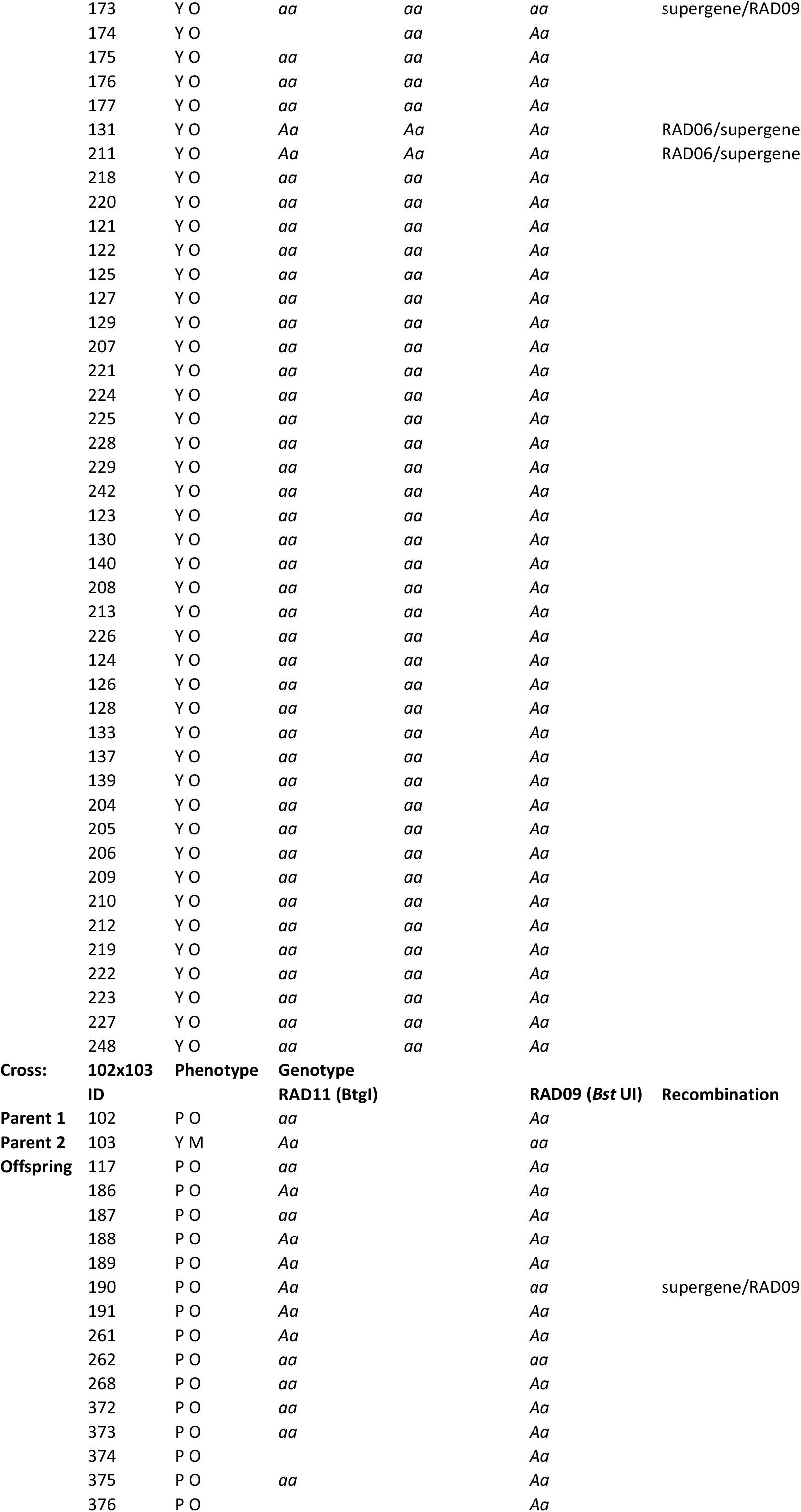
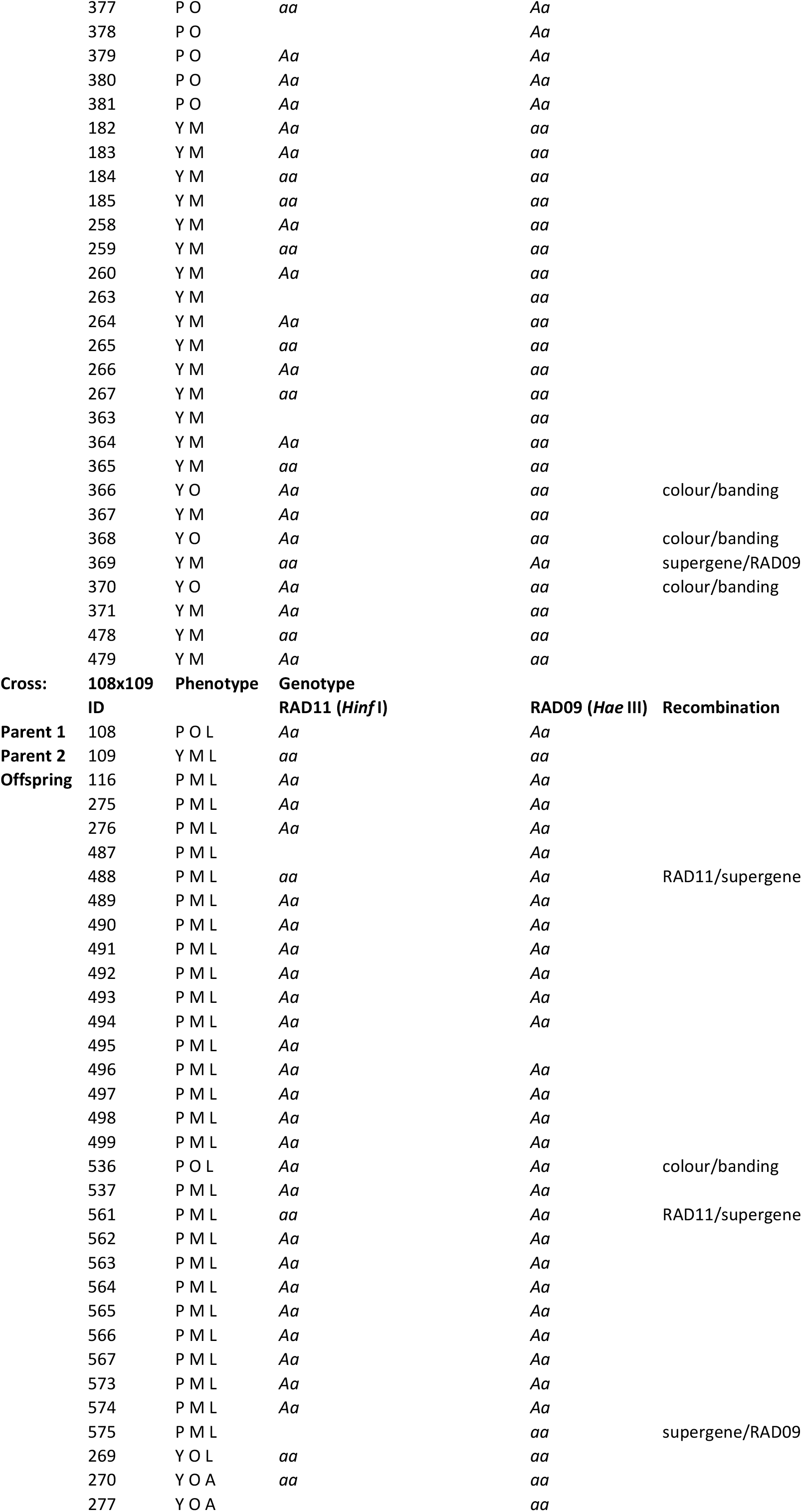
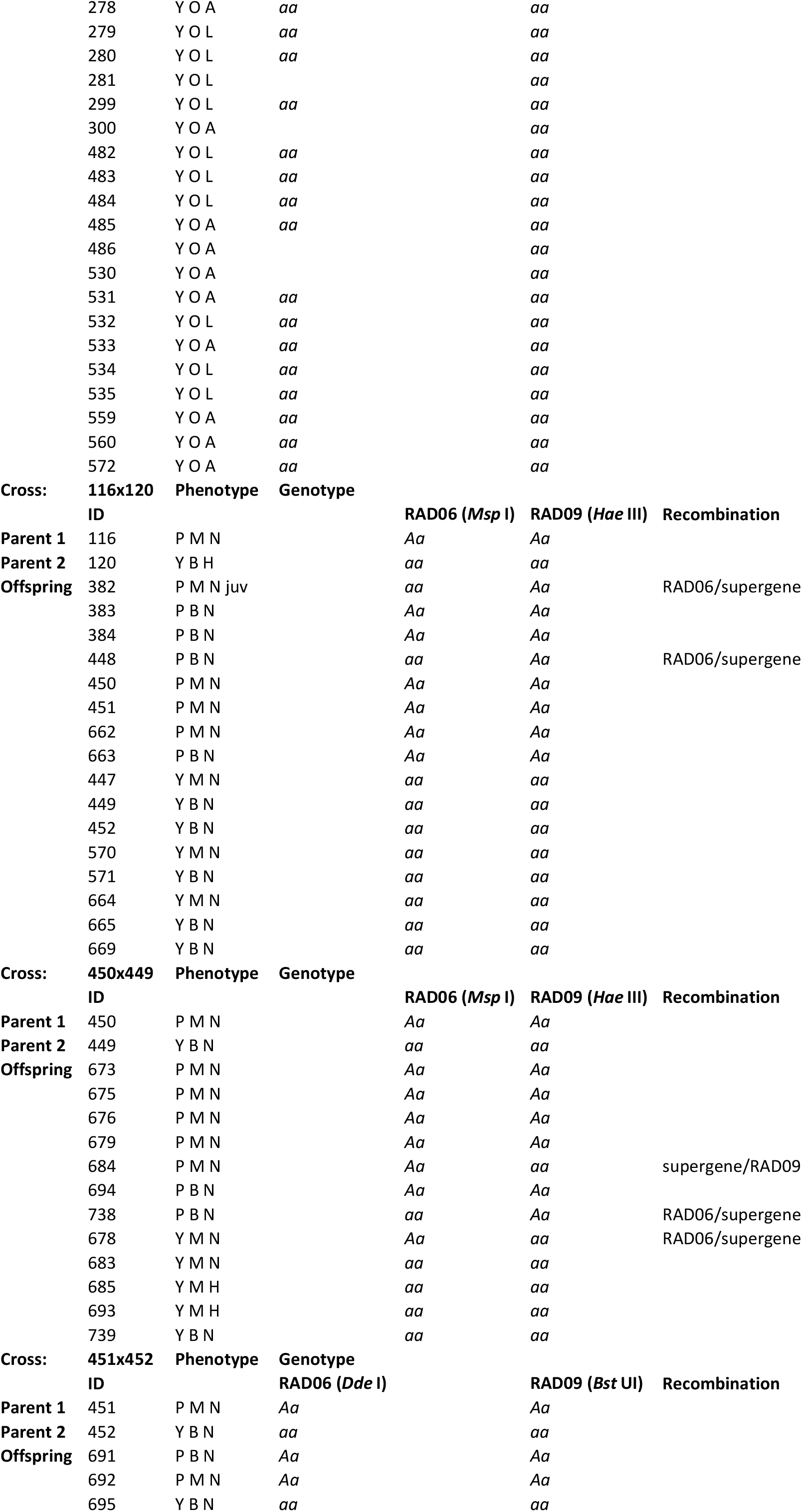
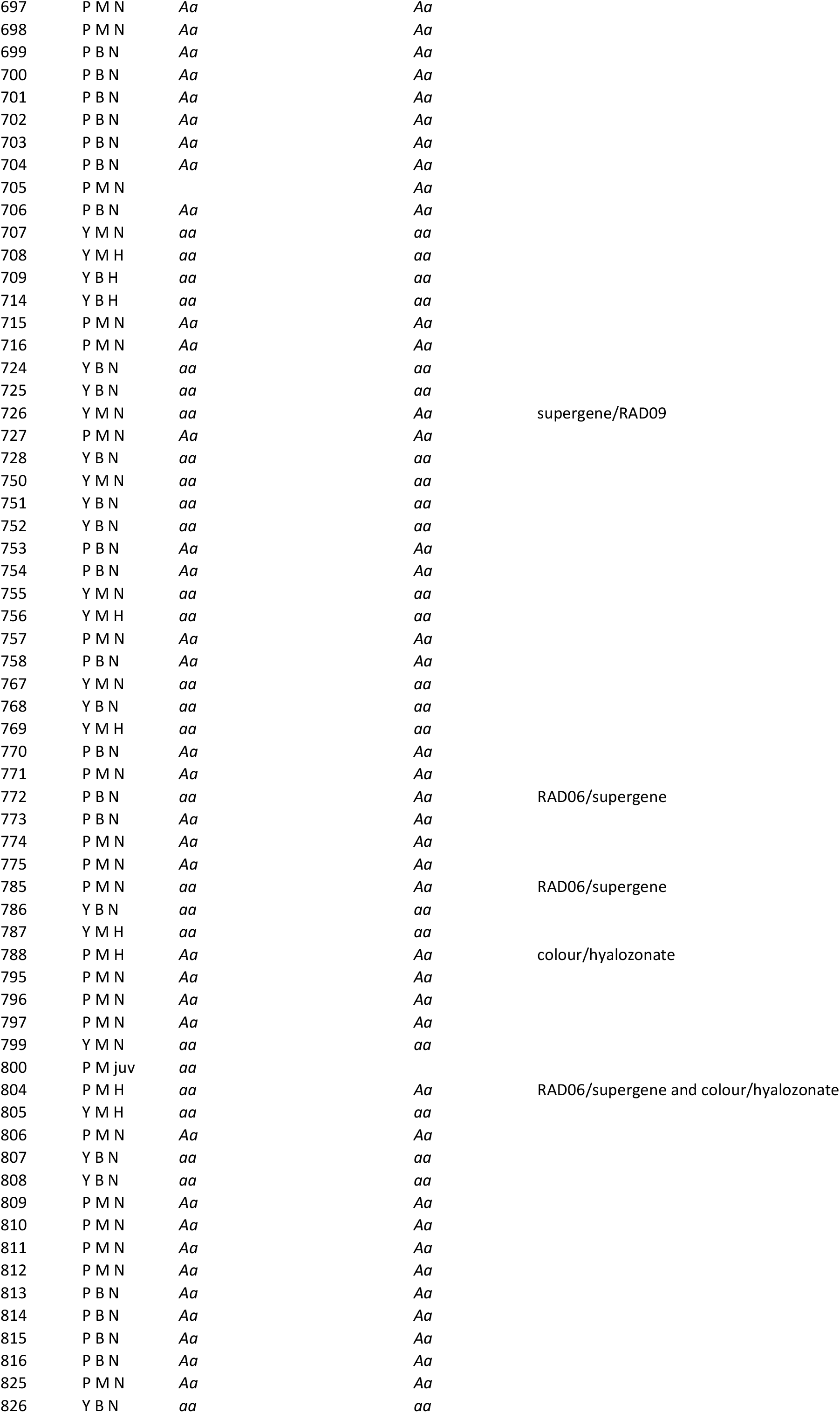
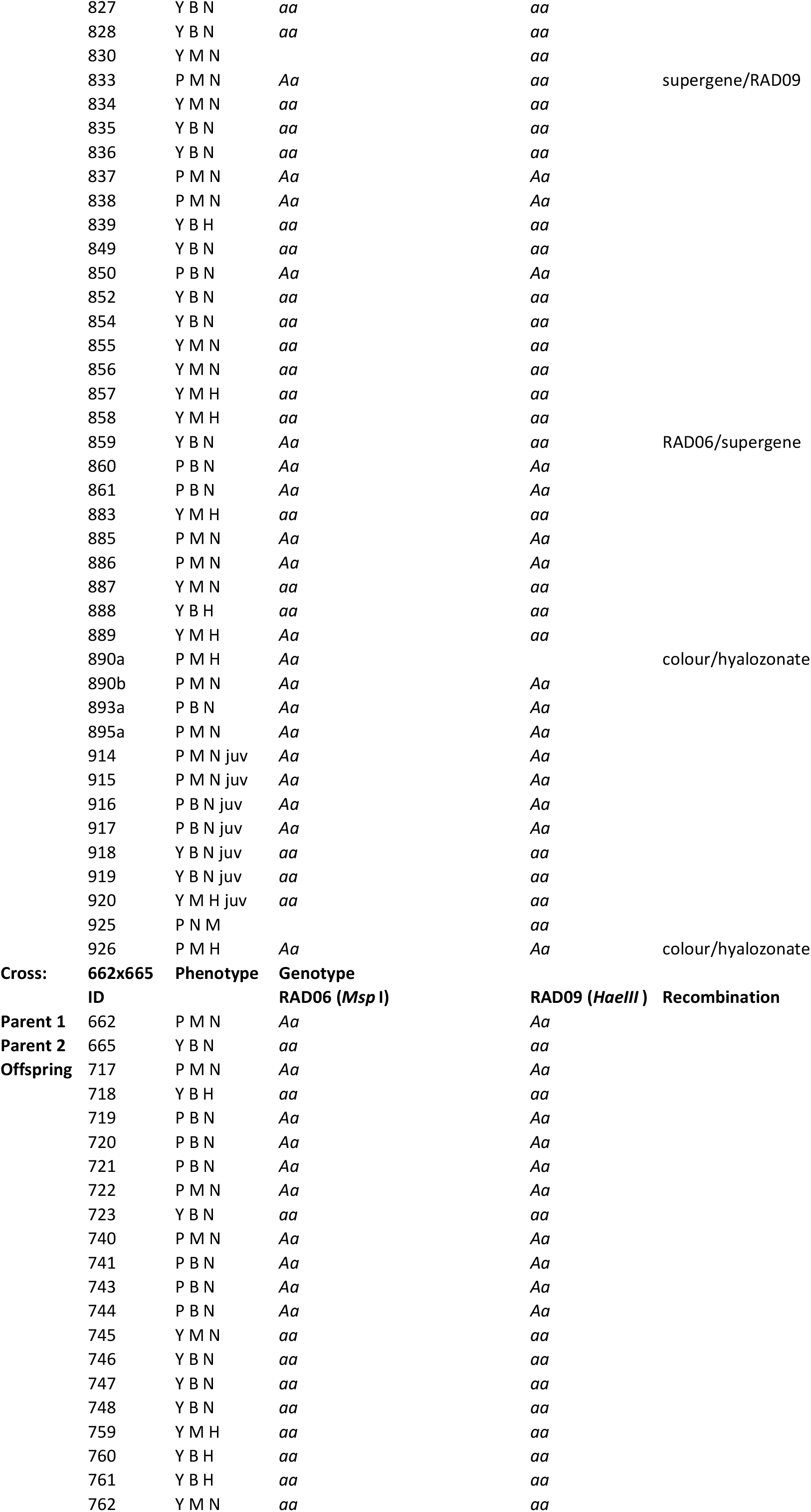
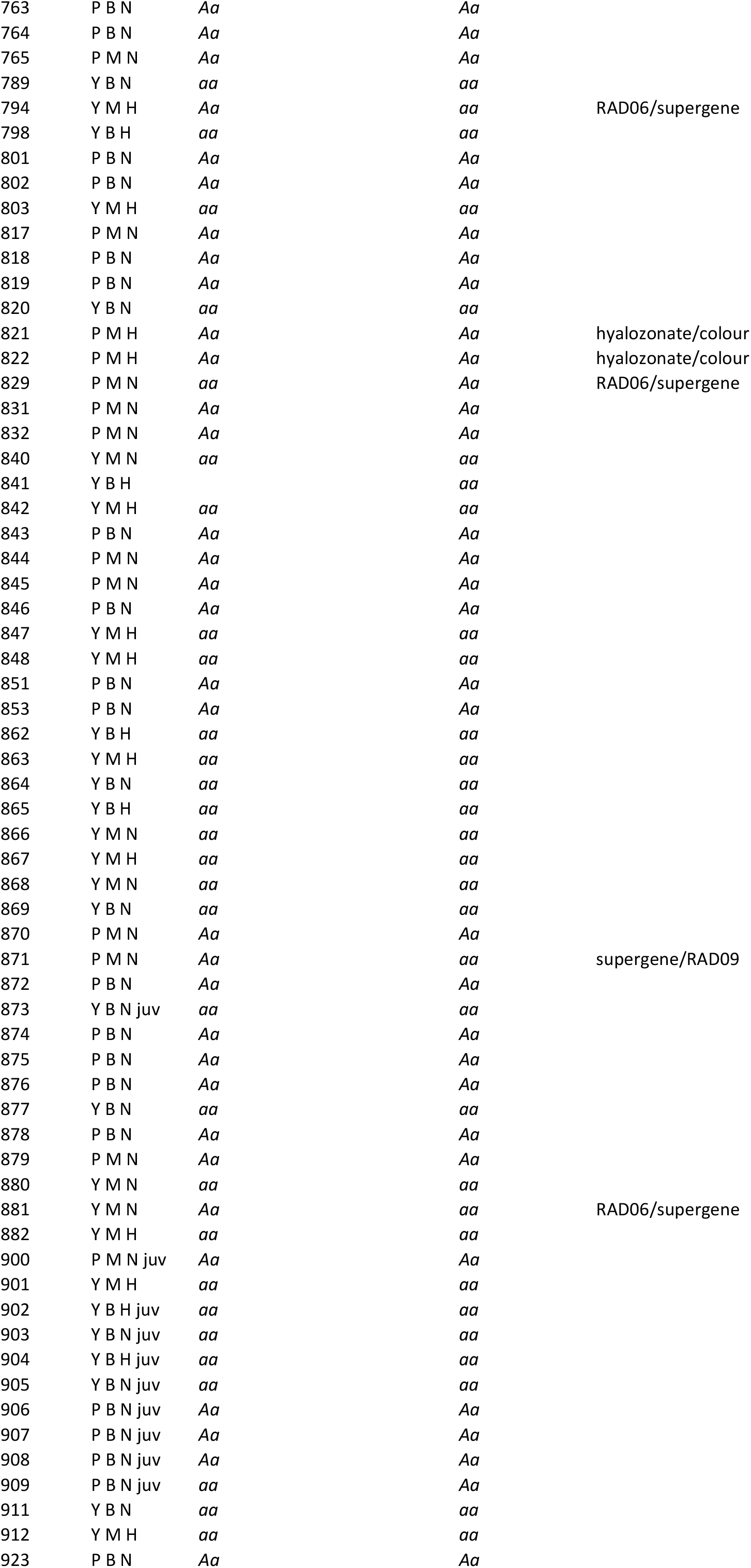
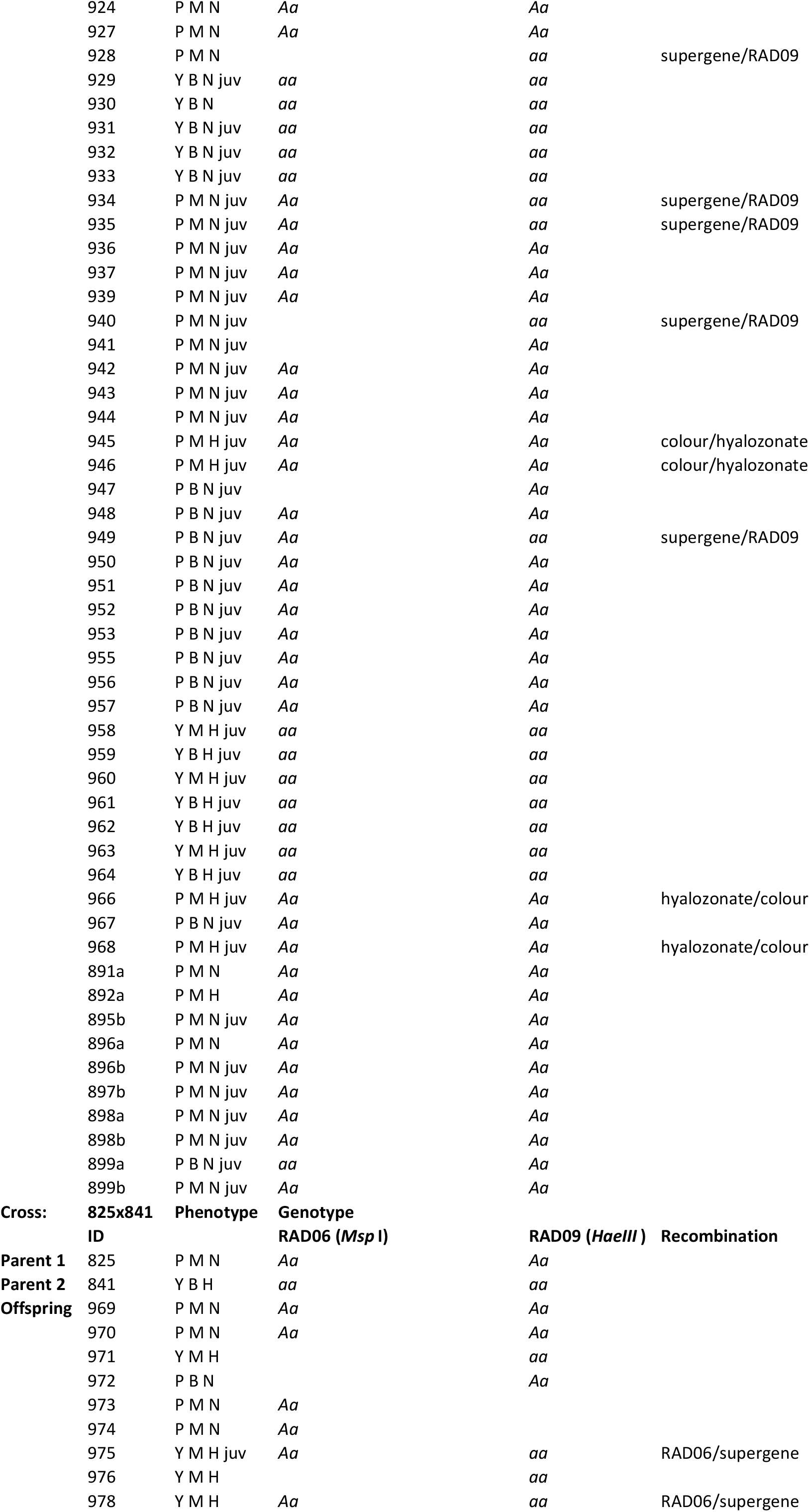
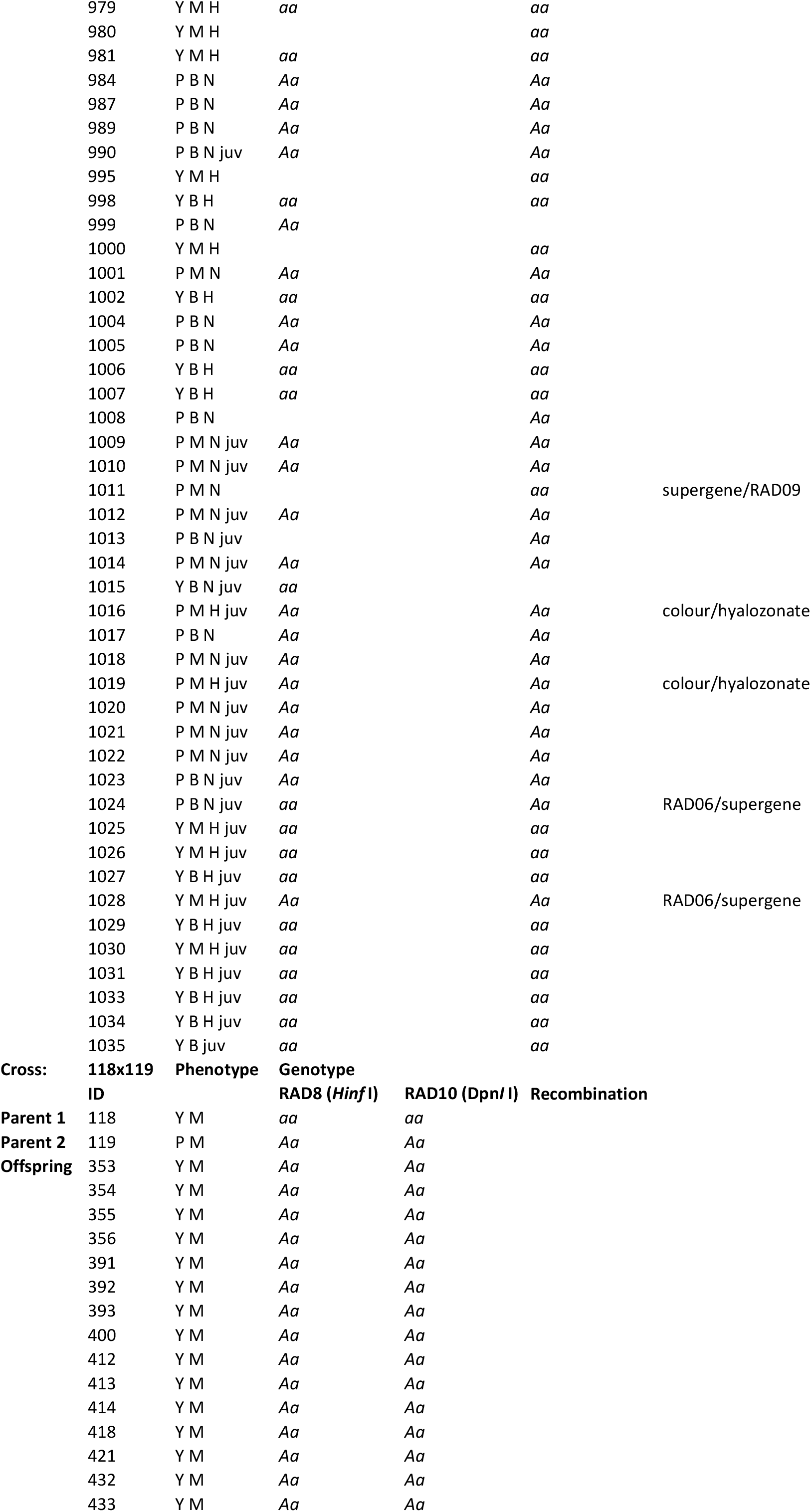
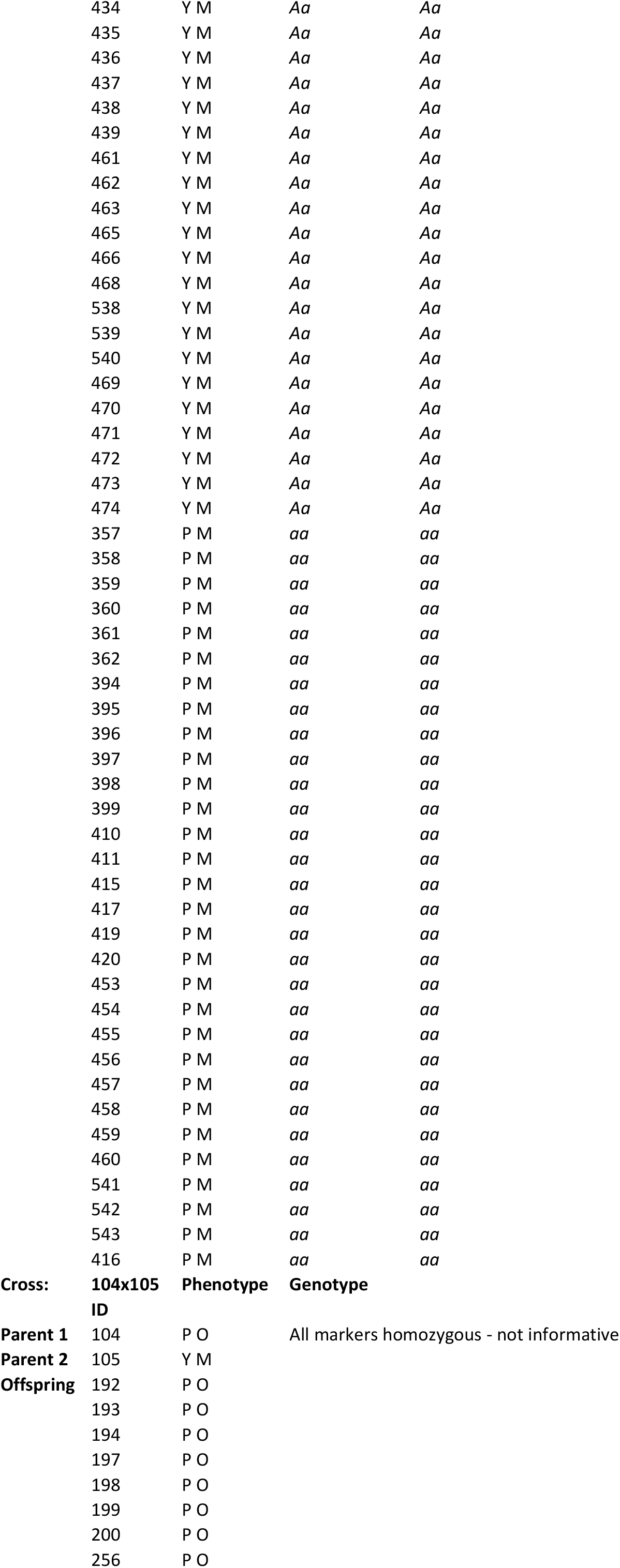
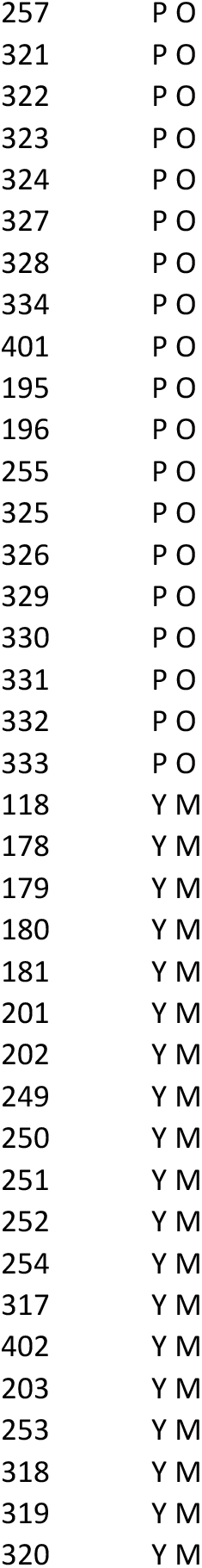
Results of genotyping for each cross, including details of restriction enzyme assay.

**Supplementary Table 2.**
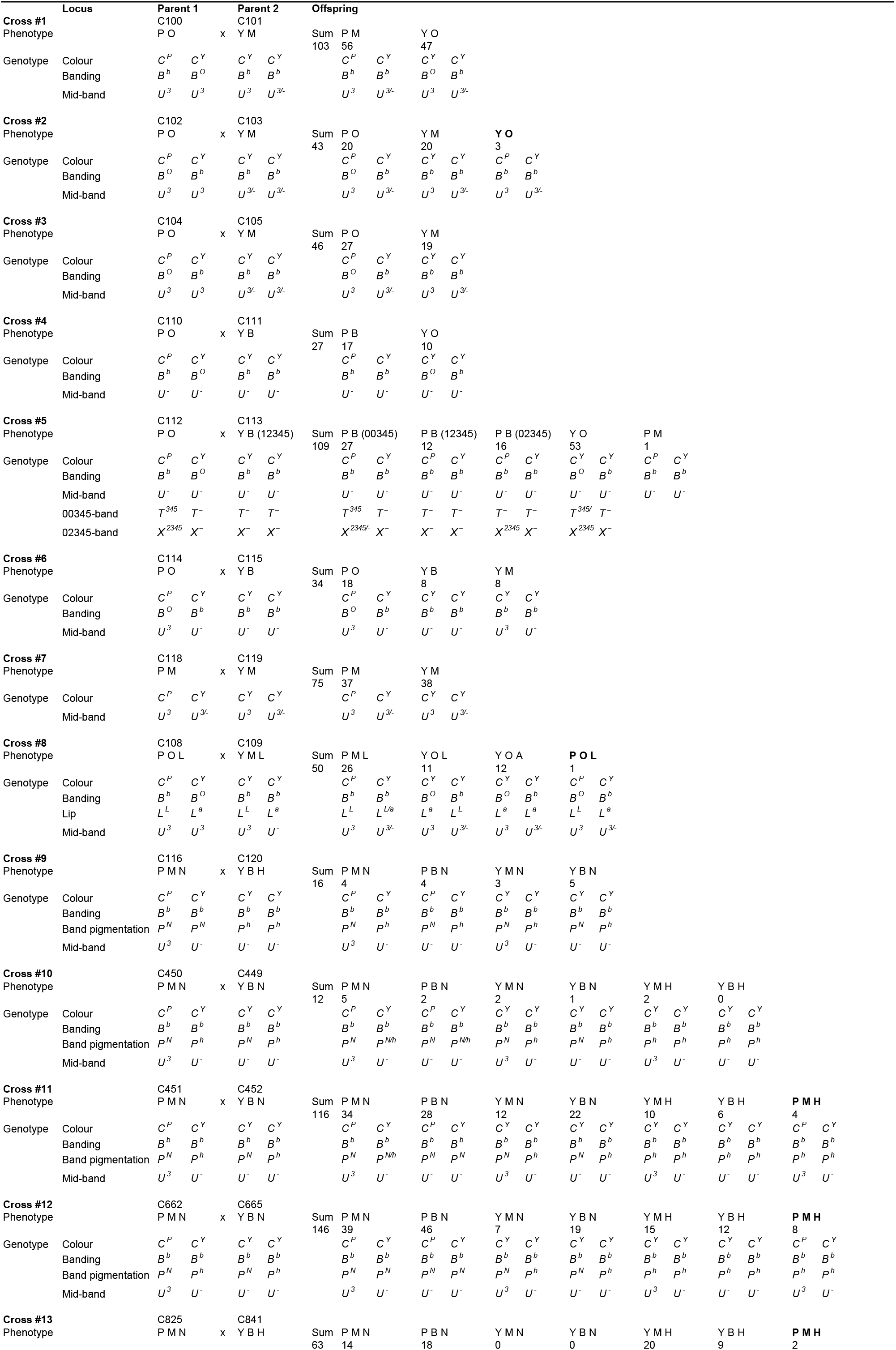
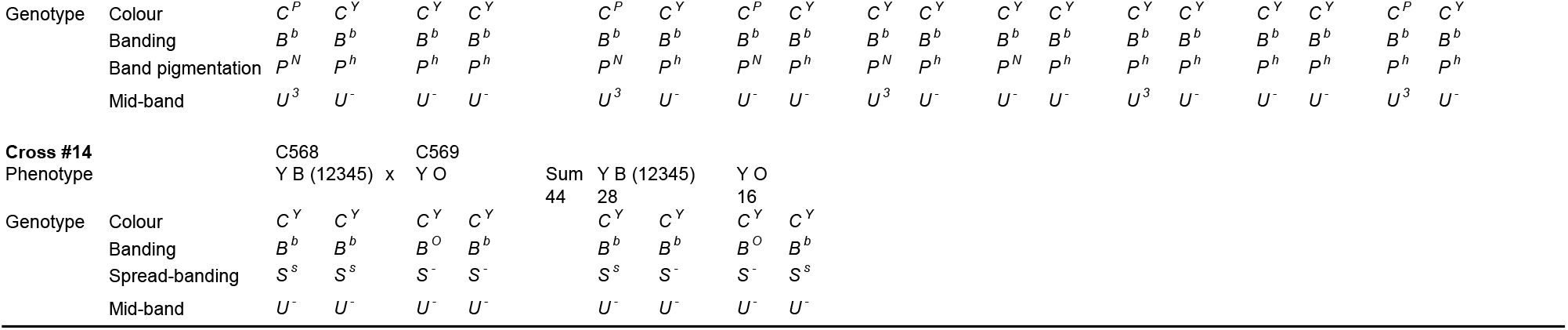
Inferred genotypes of all individuals. Shells with a phenotype that may be due to a potential recombination event in parent are shown in bold.

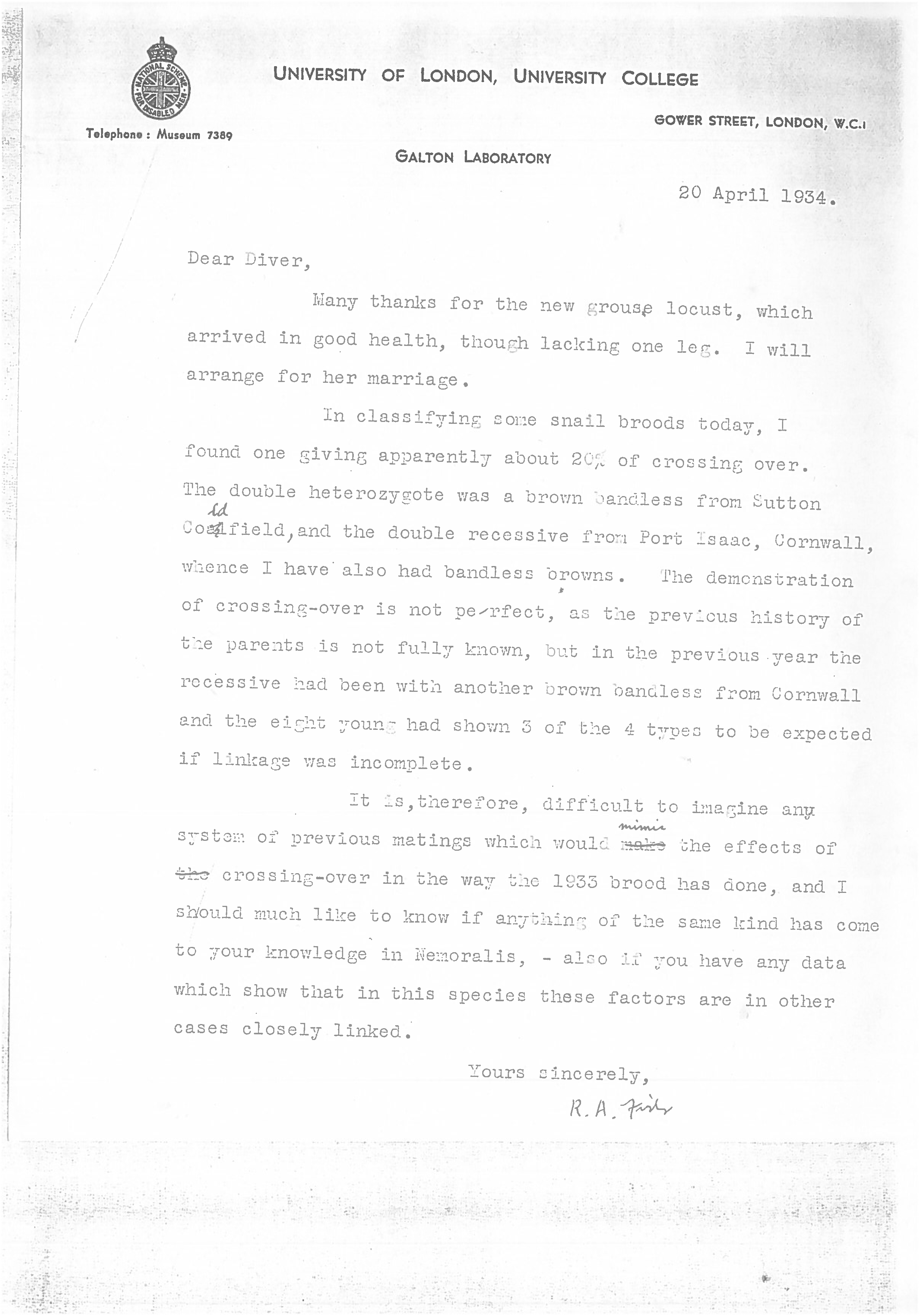
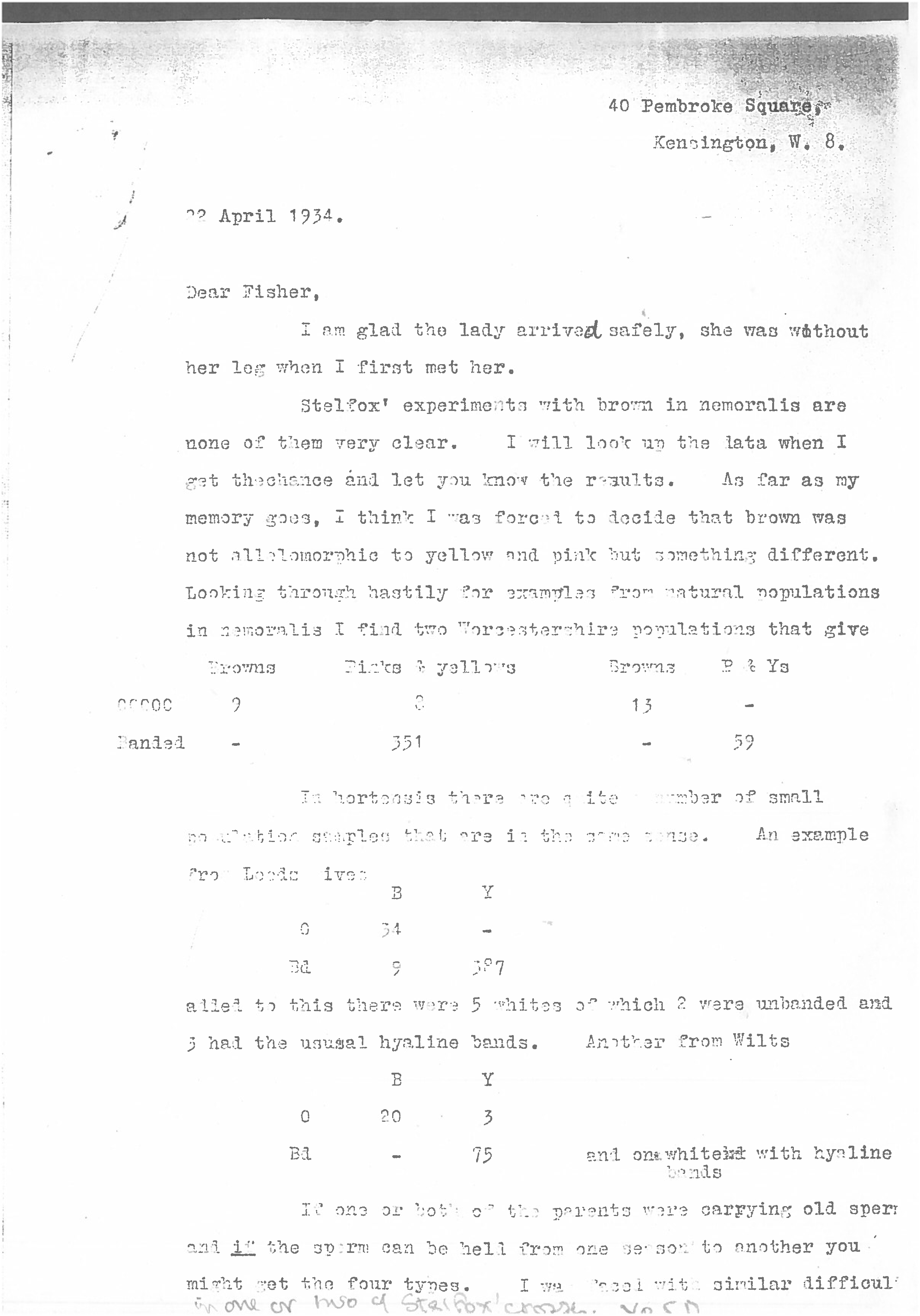
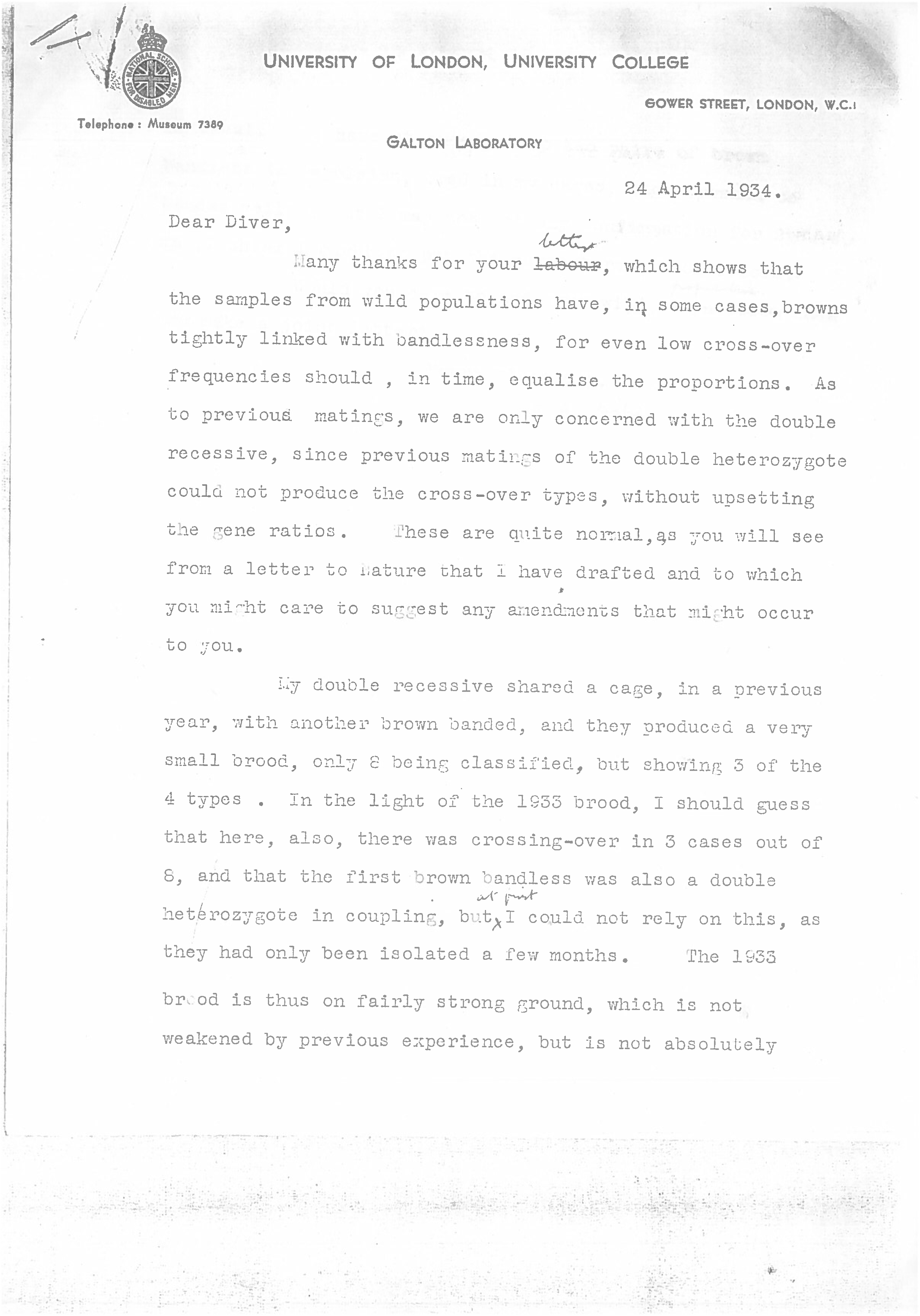
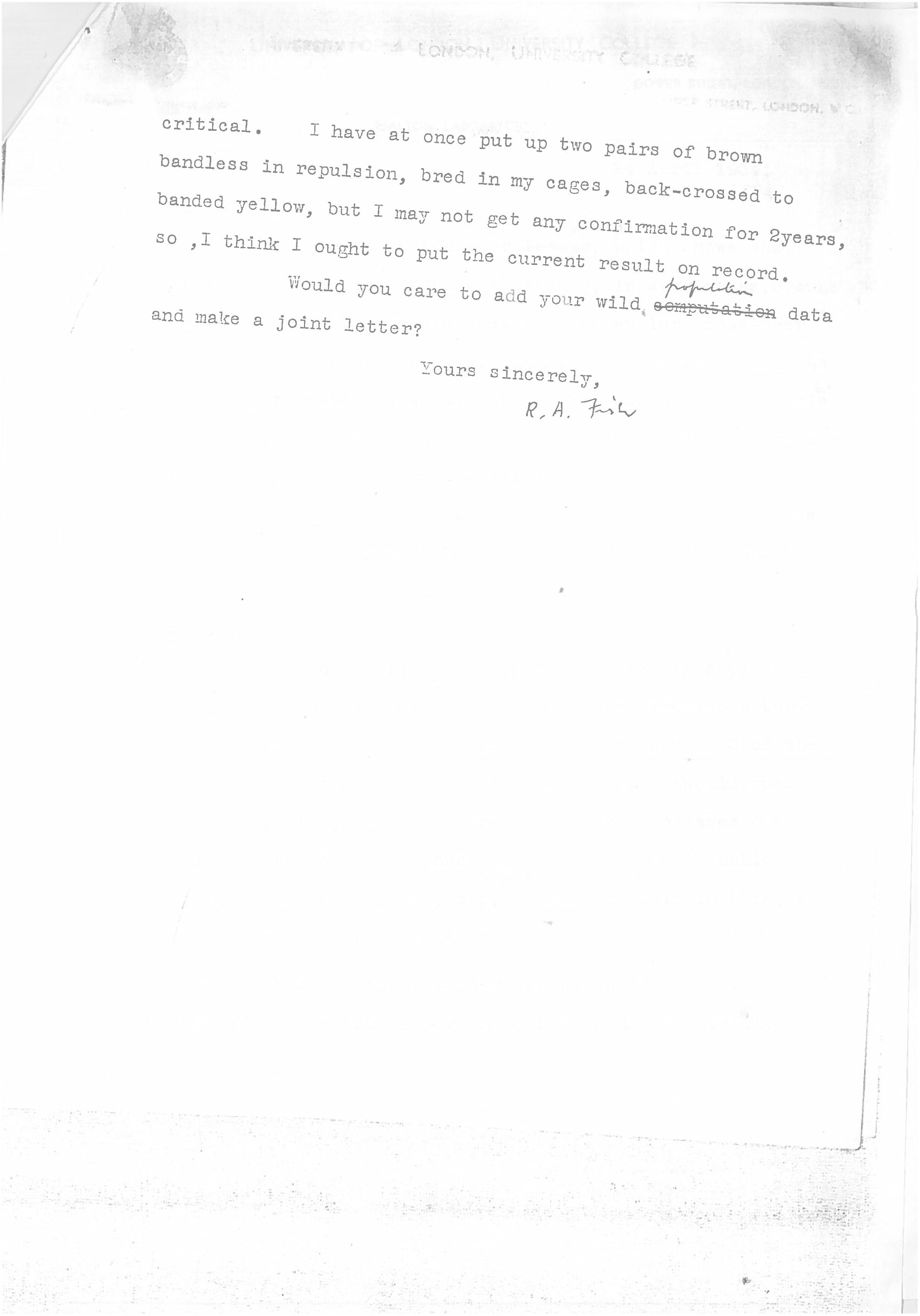
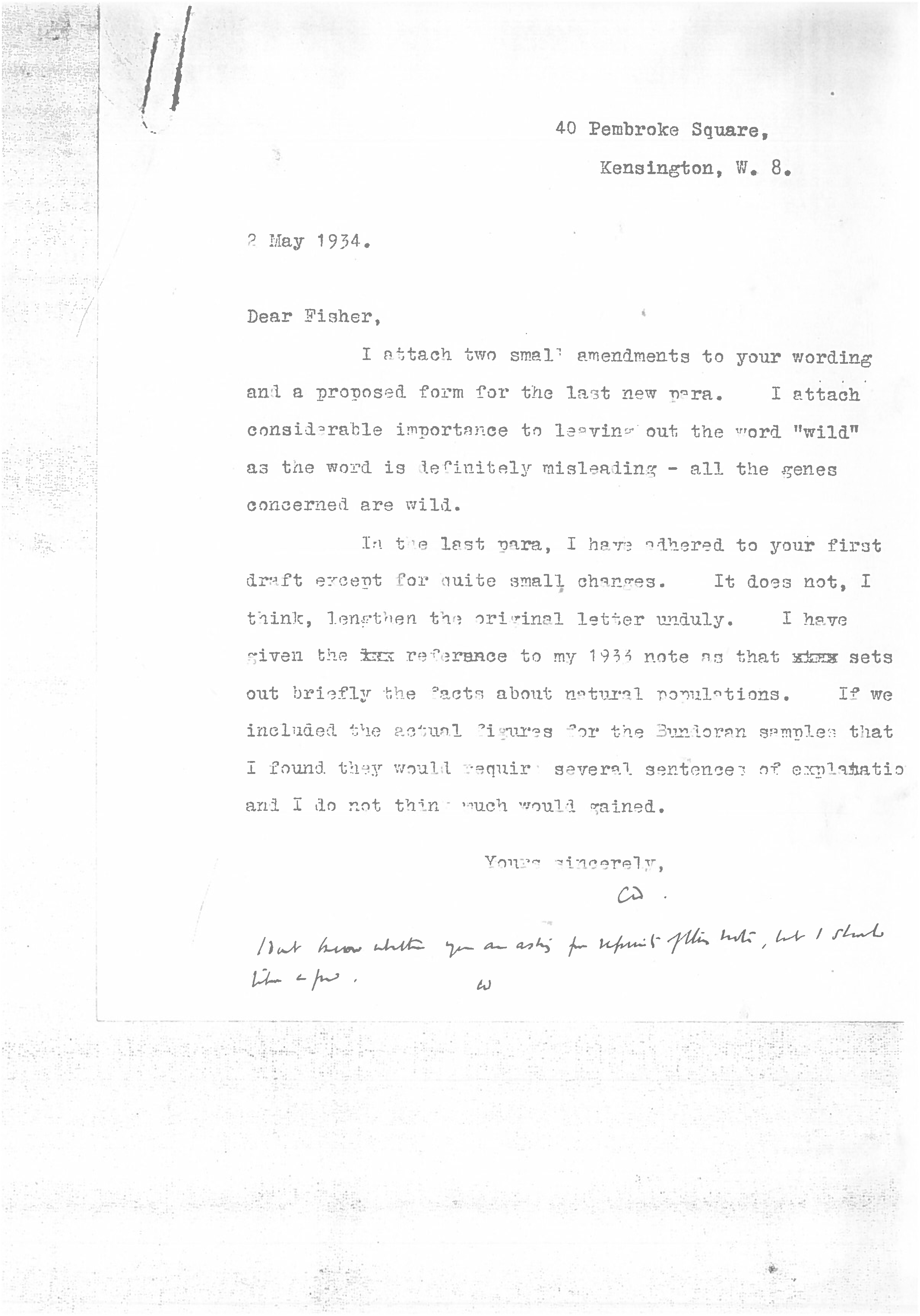
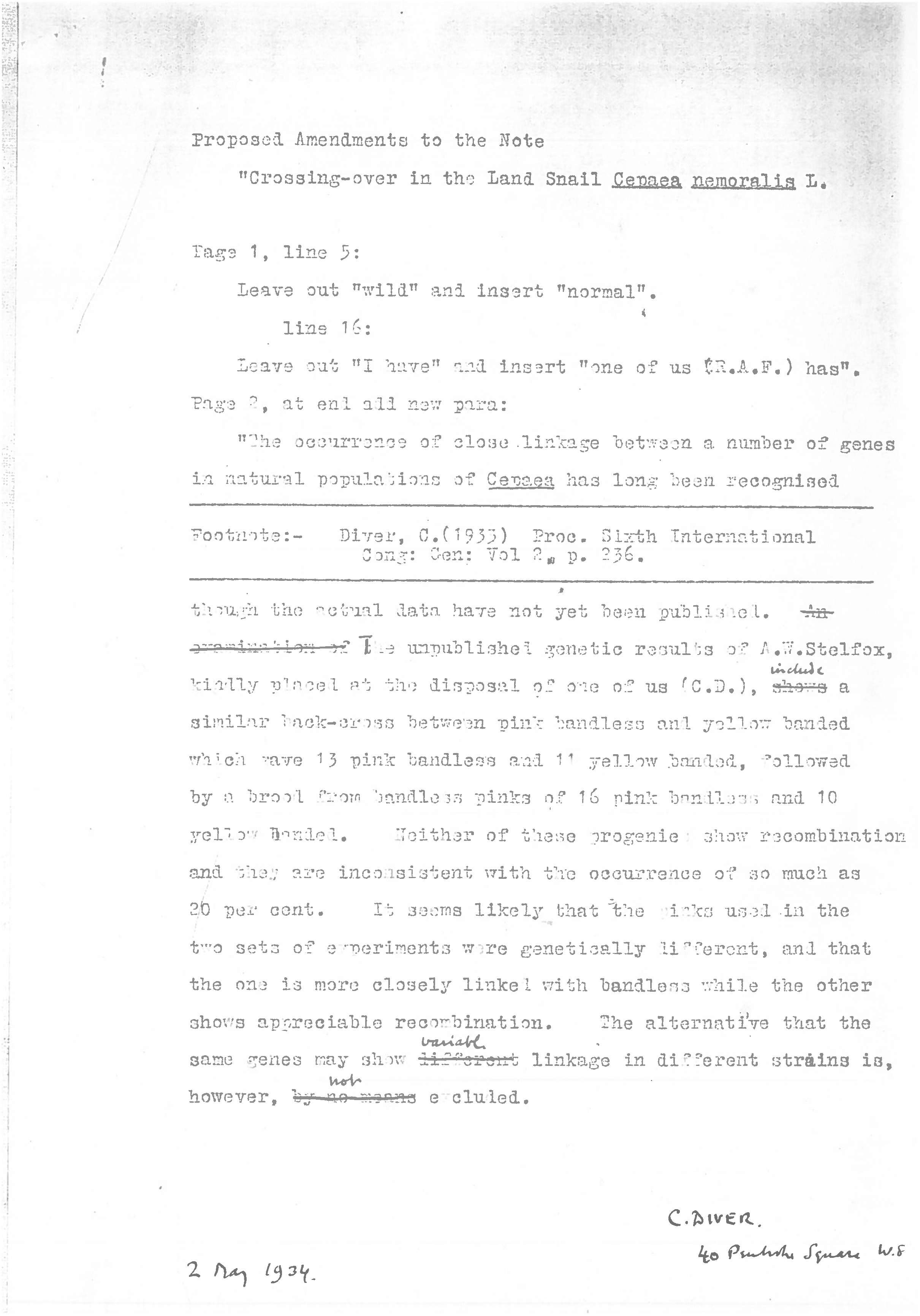
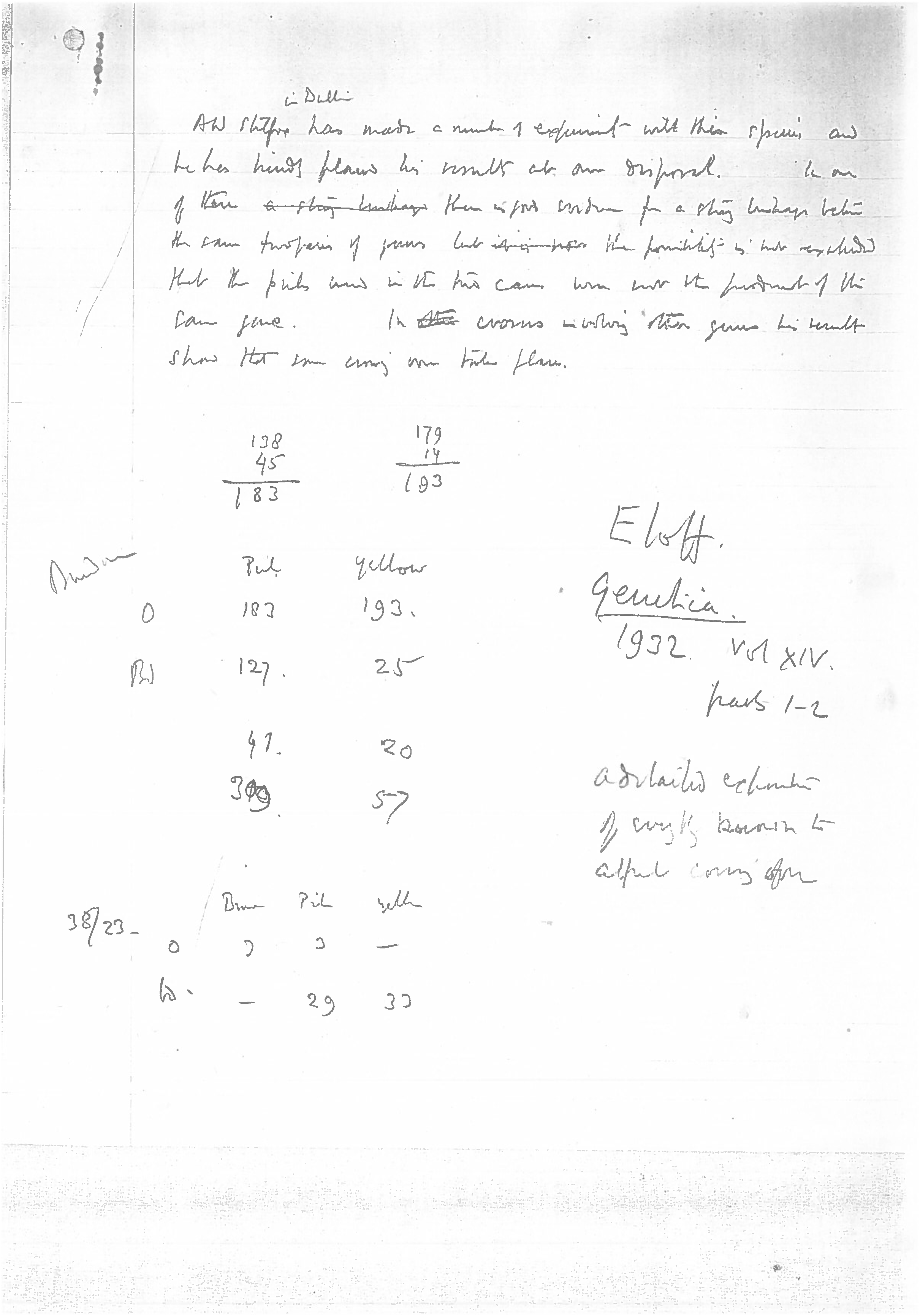
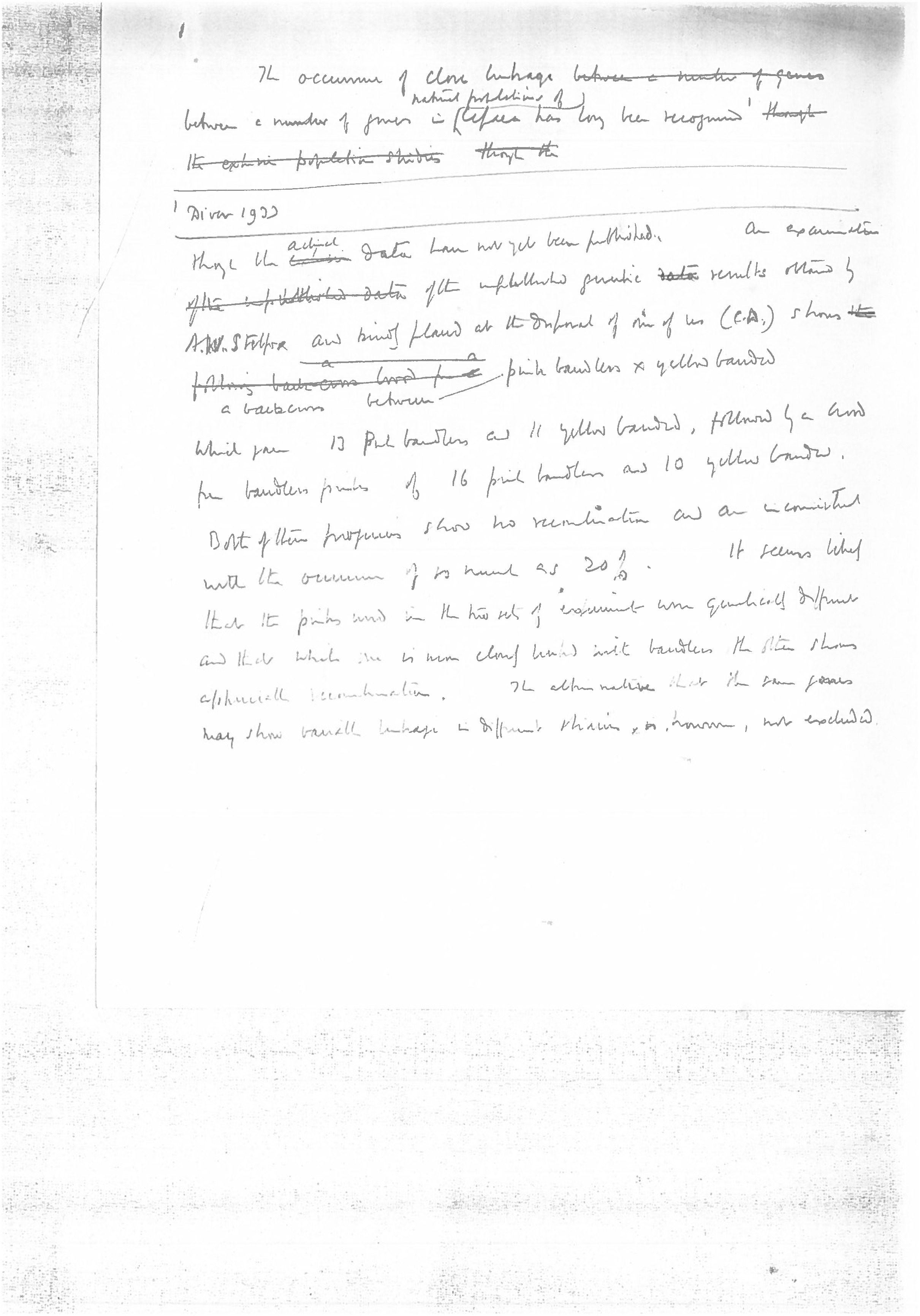
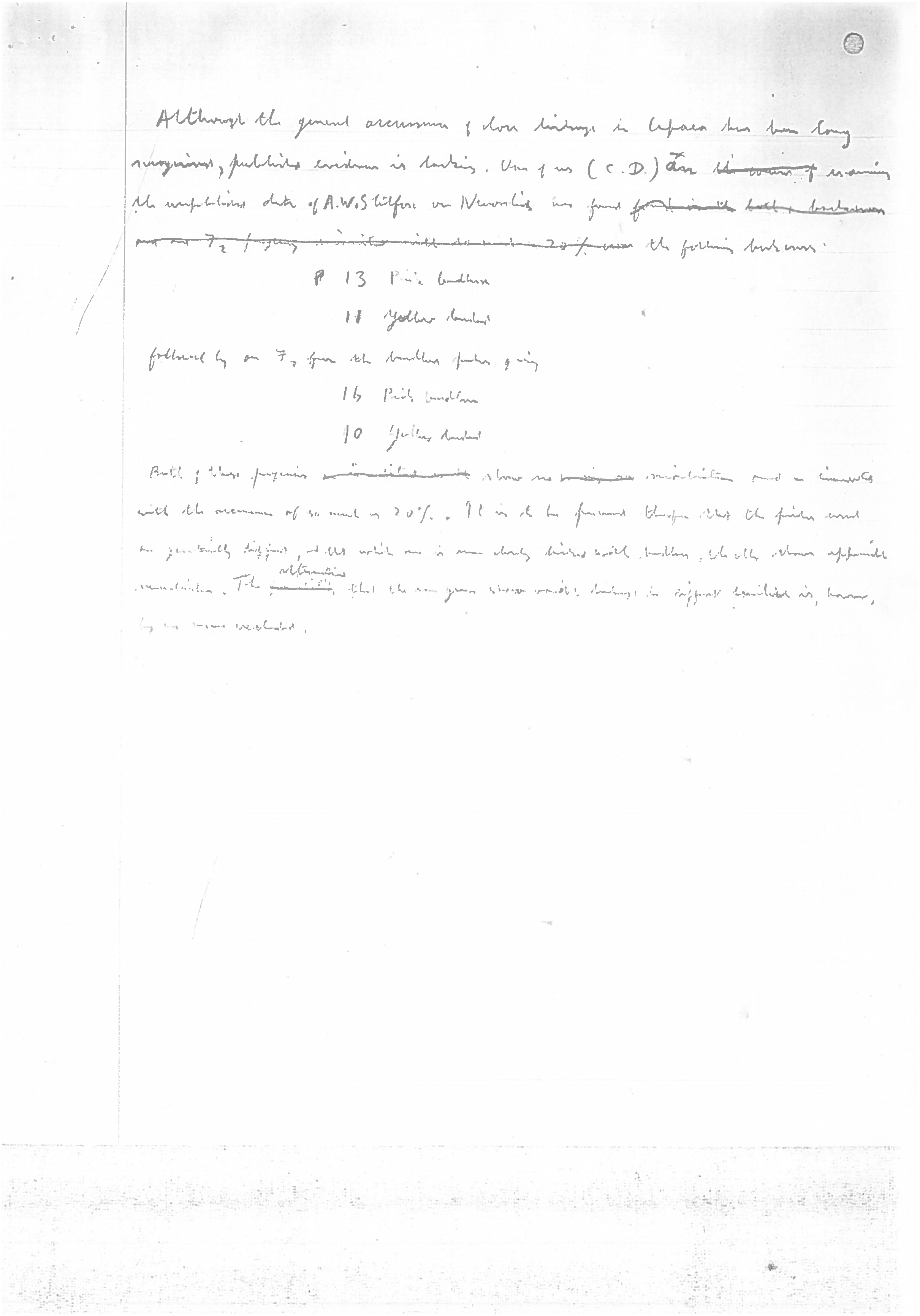
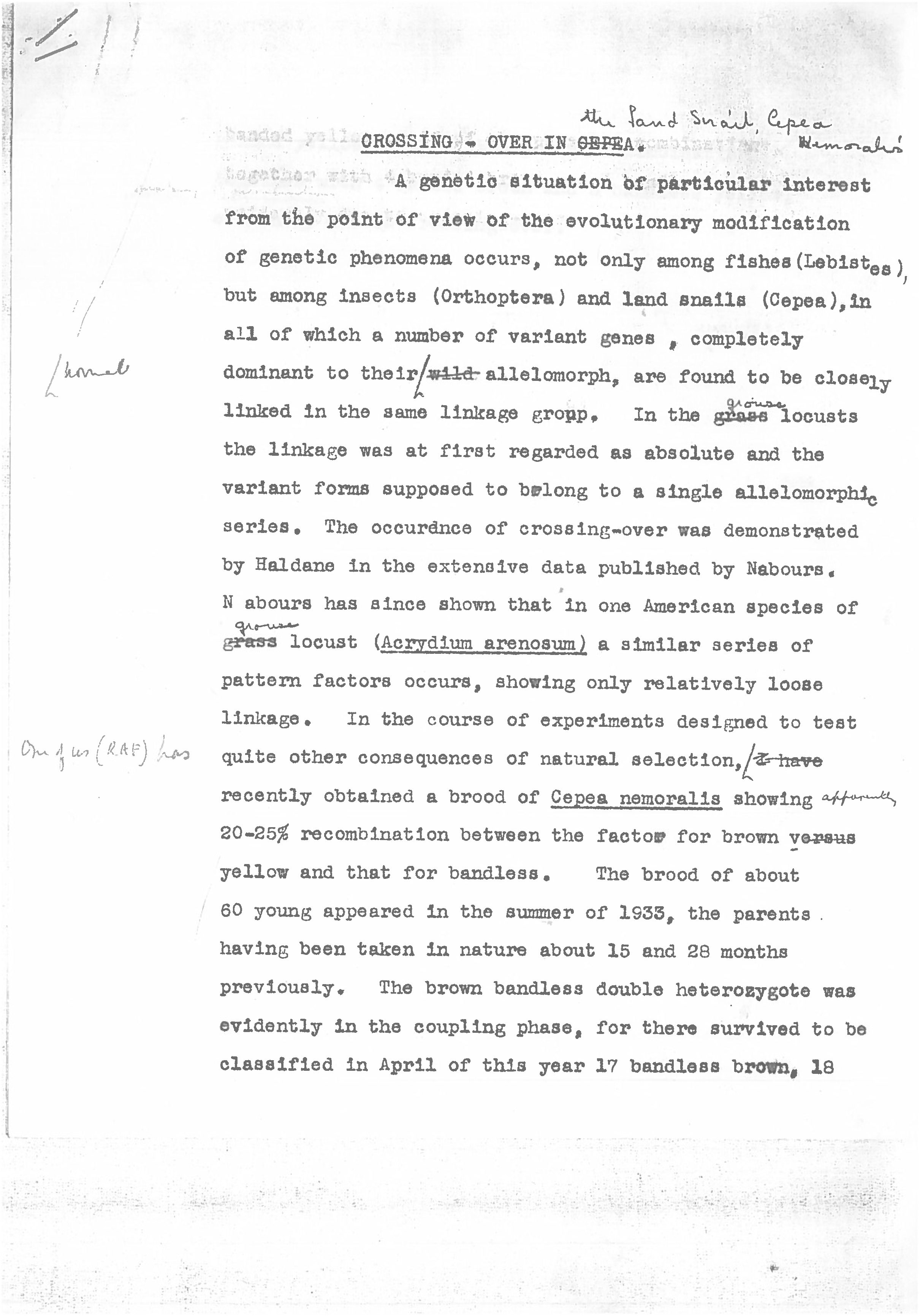
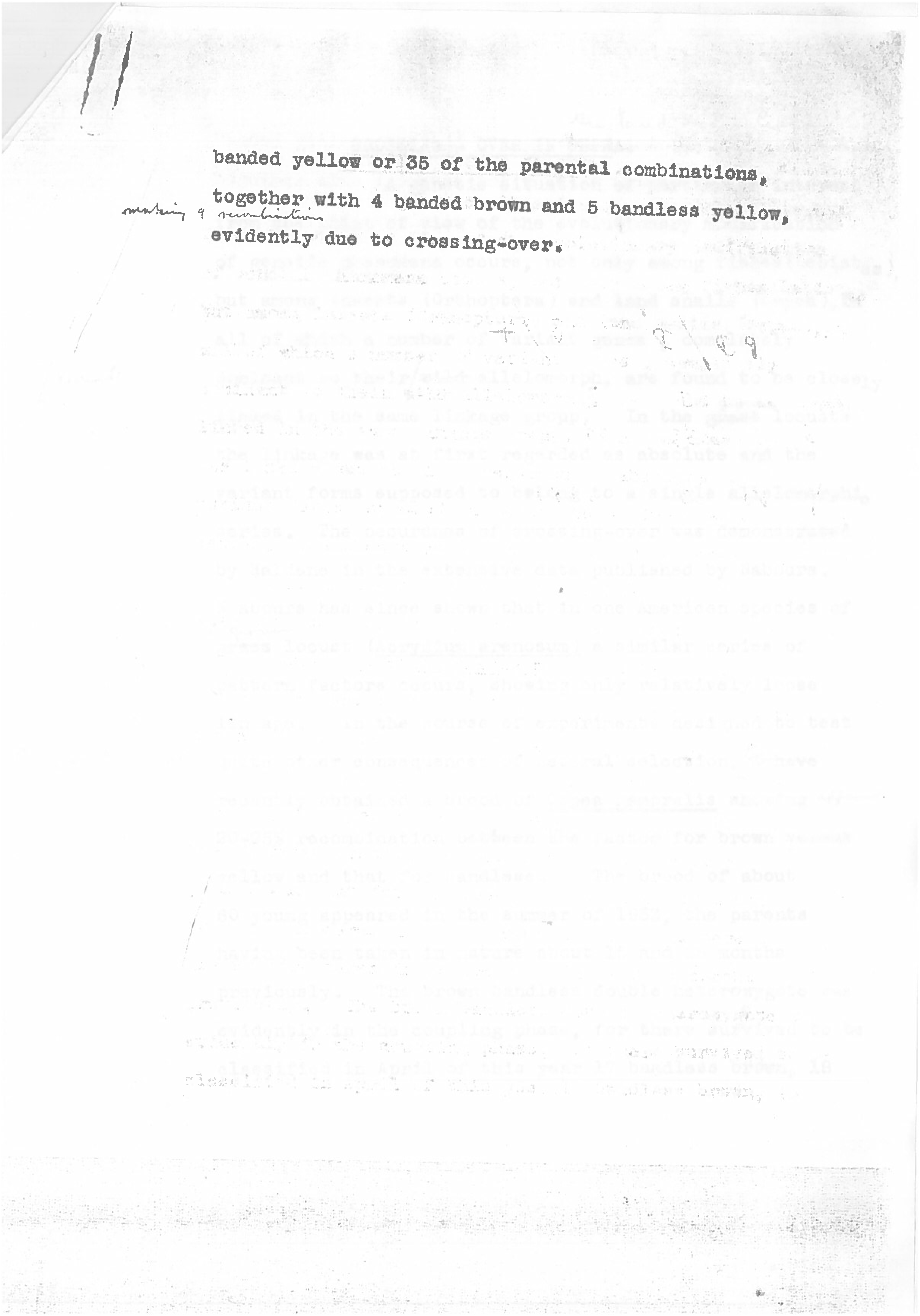

